# Hedgehog signaling drives glial cell plasticity and oncogenic reprogramming in gastroenteropancreatic neuroendocrine neoplasms

**DOI:** 10.1101/2025.09.05.674574

**Authors:** Suzann Duan, AnneLeigh B. Twer, Ateh Zinkeng, Ricky A. Sontz, Juanita L. Merchant

## Abstract

Disruption of the *Men1* locus in epithelial and endocrine tissues fails to generate the full spectrum of gastroenteropancreatic neuroendocrine tumors (GEP-NETs), raising the possibility of a potential stromal source for these cancers. Neural crest-derived glial cells were previously implicated in neuroendocrine tumors arising in the pituitary and pancreas, yet these studies lacked a clear mechanism for these events. Here, we investigated the hypothesis that *Men1*-driven Hedgehog (HH) signaling redirects the glial cell fate to give rise to neuroendocrine tumors in the gastrointestinal tract. Using archived patient specimens and The Cancer Genome Atlas, we demonstrated that GEP-NETs overexpress HH pathway components, including SHH and its cognate receptor PTCH1. We showed that patient-derived pancreatic NET tumoroids proliferate in response to SHH pathway agonists. In contrast, pharmacologic inhibition of GLI1/2, but not inhibition of SMO alone, attenuated tumoroid growth. Using two genetic mouse models, we showed that loss of *Men1* in GFAP^+^ and SOX10^+^ glial cells causes the development of pancreatic and small intestinal NETs that overexpress HH proteins. Further use of tdTomato^+^ mice demonstrated the involvement of GFAP^+^ and SOX10^+^ glial cells in these tumors. Tumoroid cultures of mouse pancreatic and small intestinal NETs recapitulated the drug response shown by patient-derived tumoroids. Lastly, *Men1*-deficient enteric glial cultures showed a glial-to-neuroendocrine transition that was alleviated upon HH inhibition, and these events were reproduced in genetic mice harboring GFAP^+^ cells with impaired primary cilia. Our study implicates the HH signaling pathway in GEP-NET development and underscores a glial cell of origin for these tumors.

**One Sentence Summary:** *Men1*-deficient glial cells reprogram along a SHH-GLI signaling axis and are a potential source for gastroenteropancreatic neuroendocrine tumors.

## INTRODUCTION

Gastroenteropancreatic neuroendocrine tumors (GEP-NETs) comprise the most common NET subtype and are rapidly increasing in prevalence, indicating a strong need for developing new approaches to detect and manage these cancers in a growing patient population (*1,2*). GEP-NETs encompass remarkably diverse neoplasms that vary in clinical presentation, mutational status, and response to therapy. Recent efforts to clarify their etiology revealed intra- and inter-tumor cell heterogeneity that was suggested to arise from reprogrammed resident endocrine cell populations (*3–5*). A deeper understanding of these reprogramming events and how they contribute to the etiology of these cancers is precluded by a dearth of preclinical models from multi-organ NETs that include pancreas and small intestine tissue sources. Modeling extra-pancreatic gastrointestinal NETs poses a distinct challenge, likely due to the relative low abundance of driver mutations in these neoplasms and reports of tumors containing a mixture of cell populations (*5–10*)*. De novo* mutations in the *Multiple Endocrine Neoplasia I* (*MEN1*) gene encoding the tumor suppressor protein Menin account for up to 40 percent of human GEP-NETs (*11,12*), yet disruption of the *Men1* locus in epithelial and endocrine tissues in mice failed to generate the full spectrum of GEP-NETs (*13–18*). These reports raised the possibility that enteric neuroendocrine cells might differentiate from a neurectoderm lineage rather than solely from endoderm-restricted progenitors. The premise that GEP-NETs can arise from neural cell progenitors challenges the current dogma and is consistent with the known neural origin of adrenal and pituitary NETs.

We and others have reported that neural crest derived glial cells, and their progenitors can acquire an endocrine phenotype leading to functional hormone secretion (*18–20*). In support of a stromal source for NETs, glial-restricted deletion of *Men1* in mice using *GFAP* and *Sox10 Cre* transgenes (*GFAP-Cre; Men1^FL/FL^* or *GFAP^ΔMen1^* and *Sox10-Cre; Men1^FL/FL^* or *Sox10^ΔMen1^*) stimulated pituitary and pancreatic NETs that coincided with loss of the glial-restricted lineage, suggesting that glial cells might undergo reprogramming upon *Men1* inactivation (*21*). Menin is a ubiquitously expressed nuclear scaffold protein that complexes with multiple transcription factors and epigenetic modifiers to regulate the expression of genes involved in cell growth and endocrine cell-specification (*22*). Among these, Menin antagonizes components of the Sonic hedgehog (SHH) signaling pathway known to direct neural cell fate patterning and the neuroendocrine phenotype (*23,24*). Consistent with this knowledge, impairment of the SHH signaling pathway in GFAP^+^ glial cells was found to reverse *Men1*-driven neuroendocrine cell hyperplasia in the *GFAP^ΔMen1^* mice (*21*). While these studies pointed to a role for the Hedgehog signaling pathway in NET development, the underlying mechanism of SHH activation in these tumors and the potential for pharmacologic intervention in preclinical models of GEP-NETs have not been thoroughly evaluated.

In the current study, we investigated the premise that GEP-NETs might originate from enteric glial cells that are instructed by SHH signaling. Using a combination of human and mouse derived tumor organoids, primary enteric glial cultures, and transgenic mouse models, we demonstrate that pancreatic and small intestinal NETs may develop from glial cells suggesting a neural crest origin for these gut tumors. We found that Menin inactivation in enteric glial cells promotes SHH-dependent reprogramming from a glial-restricted lineage to the NET phenotype. Pharmacologic and genetic inhibition of the GLI1/2 transcriptional effectors blocked GEP-NET development and underscores the potential value of targeting non-canonical SHH signaling in cancers with neuroectoderm origins. By establishing that NETs can arise from a glial cell that responds to SHH, our findings prompt a reassessment of HH inhibitors for potential therapeutic intervention in GEP-NETs that may include targets beyond inhibiting SMO.

## RESULTS

### Human GEP-NETs overexpress Hedgehog signaling components

The Hedgehog (HH) gene family encodes three distinct secreted proteins, Desert hedgehog (DHH), Indian hedgehog (IHH), and Sonic hedgehog (SHH) with critical roles in mammalian development. Of these, SHH directs the differentiation of enteric neural crest derived cells toward the neuronal lineage (*25–30*), and its aberrant activation is reported in multiple malignancies, including in GEP-NETs (*31–33*). To confirm the expression of SHH in human pancreatic (PNET) and duodenal (DNET) NETs, we collected tumor samples across three institutional centers and performed immunohistochemical (IHC) staining on sections from these tumors. The SHH expression patterns in the tumors were compared to normal tissues from non-tumor bearing patients and to normal appearing adjacent tissues in patients with respective PNETs or DNETs. In normal adjacent human pancreas, we observed the presence of SHH immunoreactive cells mainly in the endocrine islet and not in the surrounding exocrine tissue (Fig. 1A). Approximately one-half of the PNETs examined were immunoreactive for SHH and its expression was restricted to the tumor stroma or tumor cells, but not in both compartments of the same tumor (Fig. 1B). In normal duodenum, SHH expression was restricted to the base of the epithelial lumen and staining was not detected in the submucosal Brunner’s glands from which MEN1 DNETs reportedly develop (Fig. 1C) (*34*). Contrarily, in 85% of DNET cases examined, SHH expression was detected in both the preneoplastic Brunner’s glands and in tumor cells (Fig. 1D). We then evaluated the expression of the SHH receptor PTCH1 in tumors and observed robust expression in all cases of PNET cases, whereas just 50 percent of DNETs showed PTCH1 immunoreactivity (Fig. 1E).

**Figure 1.**
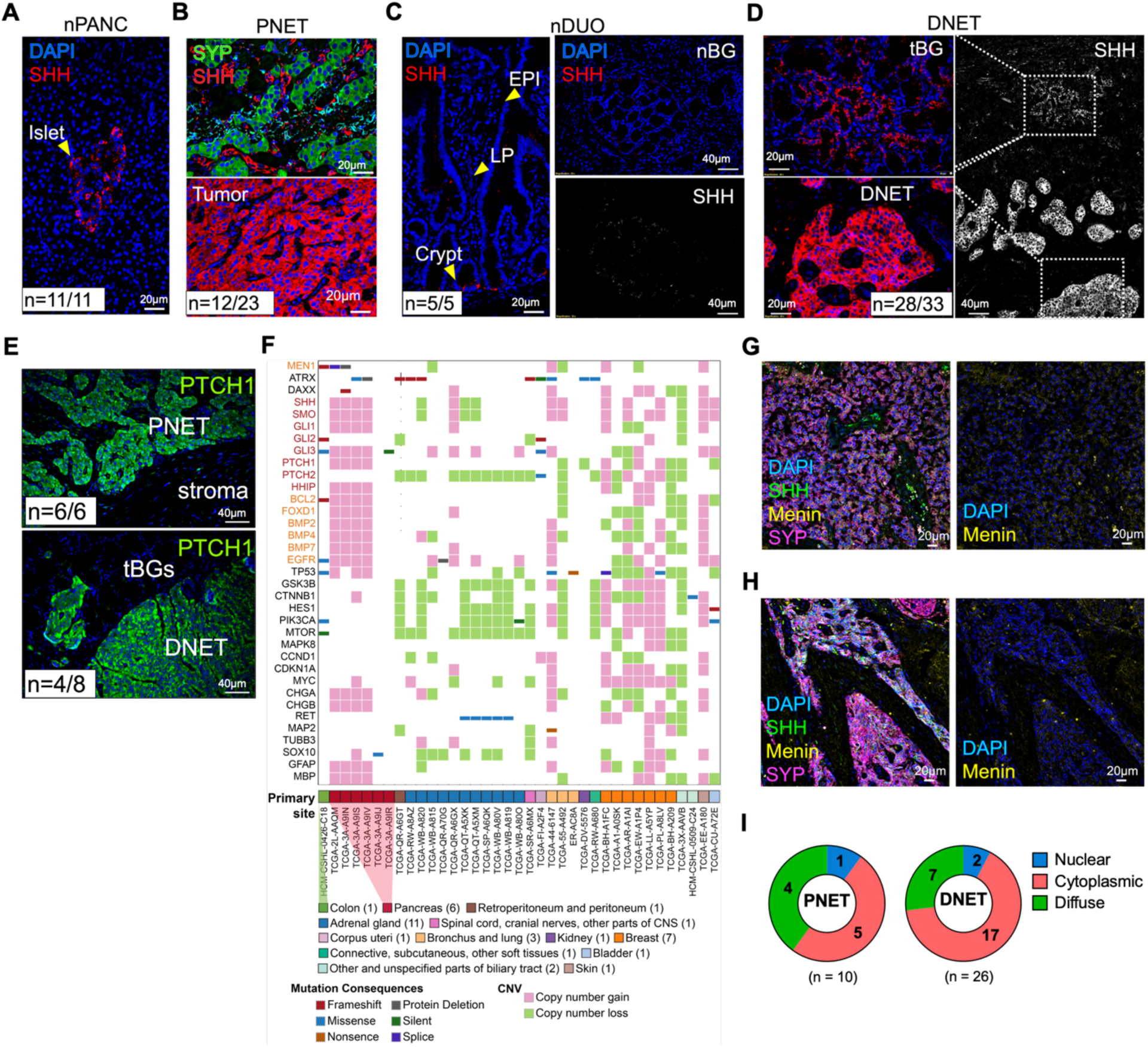
Hyperactivation of the Sonic hedgehog (SHH) signaling pathway in human pancreatic and small intestinal NETs. (**A**) Immunofluorescence (IF) images showing SHH expression (red) in normal adjacent human pancreas islet (nPANC) compared to (**B**) pancreatic NETs (PNET). SHH is expressed in the PNET microenvironment (top panel, tumor cells indicated by synaptophysin labeling in green) and by PNETs (bottom panel). (**C**) IF images of SHH staining (red) in normal duodenum (nDUO, left panel) and Brunner’s glands (nBGs, top right panel). Bottom right panel shows SHH staining (white) without DAPI overlay. Arrowheads indicate to the luminal epithelium (EPI), lamina propria (LP), and crypts. (**D**) IF images showing SHH expression (red) in transitioning BGs (tBGs) and duodenal NETs (DNET). Right panel shows the wide-field image of SHH staining (white) without DAPI overlay. (**E**) Human PNETs and DNETs were stained for the SHH receptor PTCH1 (green). For **(A–E)** n = # patients with positive staining/total number patient samples. (**F**) Heatmap of neuroendocrine tumors represented in the TCGA database displaying mutations or copy number variations (CNV) in the genes indicated on the y-axis. SHH signaling pathway genes and genes reported to interact with the SHH signaling pathway are indicated in red and orange text, respectively. (**G**) IF images of Menin expression (yellow) in human PNETs and (**H**) DNETs, showing cytoplasmic and diffuse expression patterns. Tumors were co-stained for SHH (green) and SYP (magenta) expression. (**I**) Circle plots representing the number of patient tumors exhibiting nuclear, cytoplasmic, and diffuse Menin staining.

To investigate whether increased SHH and PTCH1 expression in tumors correlated with genetic alterations in the SHH signaling pathway, we assessed The Cancer Genome Atlas (TCGA) for copy number variants (CNV) and enrichment of HH signaling genes in patients carrying a primary neuroendocrine diagnosis (TCGA-neuroendocrine). Compared to non-GI NETs, PNETs showed an enrichment in copy number gains in HH pathway genes, *e.g., SHH, SMO, PTCH1, GLI1*, and *GLI3* (Fig. 1F and Table 1), and this was associated with increased transcript expression of HH pathway genes in these tumors (fig. S1). PNETs exhibited copy number gains in several genes associated with SHH-interacting pathways, *e.g., BMP, BCL2, EGFR*, whereas neuroendocrine cases in the breast, skin, and lung tended to present with copy number gains in other growth signaling genes, *e.g., MYC, PI3K*, and *GSK3* (Fig. 1F). Consistent with prior reports that Menin antagonizes HH signaling (*23,24*), loss-of-function mutations in *MEN1* were only identified in PNETs and in one colon NET represented in the analysis. Given that human GEP-NETs were reported to express markers of glial cells (*18,35*), we explored whether genetic alterations in the HH signaling pathway are also shared by cancers with glial cell origins, including astrocytoma, glioma, and glioblastoma. Using TCGA, we examined the prevalence of CNVs in HH pathway genes for both neuroendocrine cancers and cancers with glial cell origins and found that approximately half of these cases exhibit copy number gains in *SHH, SMO*, and *GLI3* (table S1). These trends persisted across *MEN1*-mutated cases in the combined cohort, with nearly 74 percent of cases exhibiting loss of *MEN1* copy number and 47 percent with gains in the previous HH signaling genes (table S2).

**Table 1.** Prevalence of copy number variations (CNV) in hedgehog pathway genes for neuroendocrine cancers in the TCGA database.

| Gene ID | Gene Symbol | Gene Name | Cytoband | # of Cases Tested for CNVs | # of Cases with CNV Gain and (%) | # of Cases with CNV Loss and (%) |
| --- | --- | --- | --- | --- | --- | --- |
| ENSG00000106571 | GLI3 | GLI family zinc finger 3 | 7p14.1 | 215 | 35 (16.3%) | 4 (1.9%) |
| ENSG00000128602 | SMO | smoothened, frizzled class receptor | 7q32.1 | 215 | 32 (14.9%) | 12 (5.6%) |
| ENSG00000164690 | SHH | sonic hedgehog signaling molecule | 7q36.3 | 215 | 30 (14.0%) | 12 (5.6%) |
| ENSG00000111087 | GLI1 | GLI family zinc finger 1 | 12q13.3 | 215 | 22 (10.2%) | 5 (2.3%) |
| ENSG00000185920 | PTCH1 | patched 1 | 9q22.32 | 215 | 13 (6.1%) | 16 (7.4%) |
| ENSG00000117425 | PTCH2 | patched 2 | 1p34.1 | 215 | 6 (2.8%) | 113 (52.3%) |
| ENSG00000074047 | GLI2 | GLI family zinc finger 2 | 2q14.2 | 215 | 3 (1.4%) | 13 (6.1%) |
| ENSG00000133895 | MEN1 | Menin 1 | 11q13.1 | 215 | 5 (2.3%) | 35 (16.3%) |

Menin is a known epigenetic repressor of the HH signaling pathway and *MEN1* mutations have been reported in up to 40 percent of PNETs (*11,12*). In contrast, whole exome sequencing from prior studies indicate that SI-NETs generally lack mutations in *MEN1* (*7–10*). Given the lower abundance of genetic *MEN1* mutations in SI-NETs compared to PNETs, we next investigated whether altered subcellular localization of Menin could contribute to SHH overexpression in these tumors. We evaluated the expression pattern of Menin protein by performing IHC staining on our original cohort of PNETs and DNETs. We observed a striking loss of nuclear Menin expression across both tumor types, with only eight percent of the tumors retaining distinct nuclear Menin expression (Fig. 1, G–I). Collectively, reduced nuclear Menin expression, either in the presence or absence of *MEN1* mutation, coincided with upregulated SHH signaling in human PNETs and SI-NETs.

### SHH regulates PNET growth via GLI1/2 signaling

Classical activation of HH signaling occurs when the SHH ligand binds its cognate receptor PTCH1 and activates Smoothened (SMO) at the primary cilium, a nutrient sensing organelle expressed on stromal cells including enteric glia and select endocrine cell types (*36*). Aberrant SHH activation, either through canonical SMO signal transduction or non-canonical activation of the transcriptional effectors GLI1/2, promotes tumor cell growth in basal cell carcinoma (BCC), medulloblastoma, and GEP-NETs (*23,33,37–39*). As prior studies in NETs were limited to the use of monoclonal cell lines, we sought to develop an improved preclinical model and test these events in primary tumor organoids (tumoroids) derived from surgically resected patient PNETs. As expected, patient-derived tumoroids strongly expressed the neuroendocrine marker CHGA, in addition to SHH and PTCH1 (Fig. 2A). We then tested whether PNET tumoroids respond to treatment with SHH and vismodegib, an FDA-approved inhibitor of SMO. In the presence of SHH, PNET tumoroids increased their expression of neuroendocrine transcripts (*e.g., SYP* and *CHGA*), whereas treatment with vismodegib reduced *CHGA* mRNA levels. Given that SHH signaling regulates the cell fate of neural progenitors to acquire a neural or glial cell phenotype, we measured the expression of *GFAP* as a classical marker of the glial-restricted lineage. Consistent with its role in reprogramming the glial lineage, SHH treatment downregulated the expression of *GFAP* in PNET tumoroids, whereas treatment with vismodegib increased *GFAP* expression (Fig. 2B).

**Figure 2.**
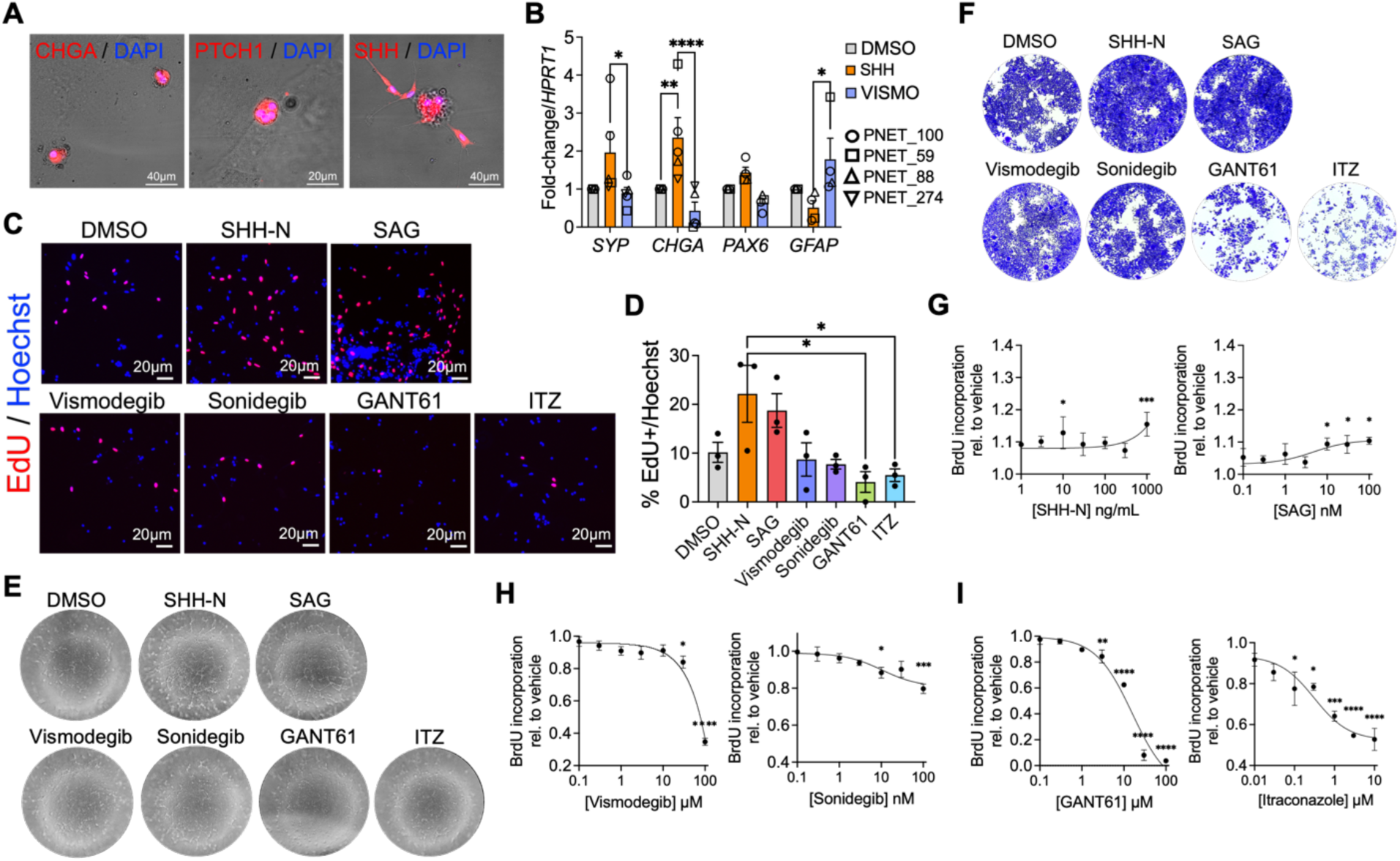
Patient-derived PNET tumoroids express HH signaling proteins and respond to HH pathway activation and inhibition. (**A**) Combined fluorescence and phase-contrast images of 21-day-old human PNET tumoroids stained for chromogranin A (CHGA) and HH proteins (PTCH1, SHH). (**B**) Patient-derived PNET tumoroids were exposed to recombinant human SHH N-terminal peptide (100 ng/mL) or the SMO inhibitor vismodegib (VISMO, 20 µM) for 72 h and changes in mRNA expression were analyzed by RT-qPCR. mRNA changes were normalized to *HPRT1* expression and DMSO vehicle control. (n = 4 unique patient lines). * = *p* < 0.05, ** = *p* <0.01, **** = *p* < 0.0001 by Two-way ANOVA with Sidak post-test. (**C**) EdU labeling showing proliferation of dissociated PNET tumoroids following 5-day exposure to: the HH agonists SHH-N (100 ng/mL) and SAG (10 nM); inhibitors of the canonical HH signaling pathway vismodegib (20 µM) and sonidegib (10 nM); or inhibitors of the GLI1/2 effectors GANT61 (10 µM) and itraconazole (ITZ, 1 µM). (**D**) Quantitation of the percentage of EdU-positive PNET tumor cells following 5-day treatment. (n = 3 replicates from one patient tumoroid line; patient ID PNET_88). * = *p* < 0.05, by One-way ANOVA with Tukey post-test. (**E**) Wide-field phase contrast images of dissociated PNET tumor cells derived from Patient ID PNET_274 following 5-day treatment. (**F**) Crystal violet staining of human BON-1 PNET cells after 48 h treatment. (**G**) BrdU incorporation in BON-1 cells after 48 h treatment with HH agonists SHH-N and SAG, (**H**) SMO inhibitors vismodegib and sonidegib, and (**I**) inhibitors of GLI1/2 signaling. * = *p* < 0.05, ** = *p* <0.01, *** = *p* < 0.001, **** = *p* < 0.0001 by One-way ANOVA with Dunnett post-test.

Next, we evaluated how modulating SHH pathway activation regulated proliferation of PNET tumoroid cells. We observed a two-fold increase in PNET tumoroid proliferation following treatment with SHH and SAG, a potent <u>S</u>MO <u>ag</u>onist, as demonstrated by increased 5-ethynyl-2’-deoxyuridine (EdU) incorporation by tumor cells (Fig. 2C). Treatment with the SMO inhibitors vismodegib and sonidegib did not affect proliferation, however inhibition of the GLI1/2 transcriptional effectors using either GANT61 or itraconazole, a pan inhibitor of SMO and GLI1/2, led to a two-fold reduction in EdU labeling (Fig. 2D). These observations coincided with increased neural outgrowth in a second patient-derived PNET line that failed to generate long term tumoroids (Fig. 2E). We further validated these findings by assessing proliferation of a monoclonal PNET cell line (human BON-1 cells) treated with these pharmacological modulators (Fig. 2F). Consistent with the response in patient-derived tumoroids, SHH and SAG treatment increased the proliferation of BON-1 cells in a dose-dependent manner (Fig. 2G). BON-1 cells were generally less sensitive to SMO inhibition (Fig. 2H), whereas direct inhibition of GLI1/2 resulted in significant growth suppression (Fig. 2I). Thus, SHH pathway activation stimulated PNET growth in a humanized preclinical model and pharmacologic inhibition of the non-canonical HH signaling pathway effectively blocked SHH-mediated tumor cell growth.

### Loss of *Men1* in GFAP^+^ and SOX10^+^ glial cells drives the development of HH-expressing GEP-NETs

Given that human GEP-NETs overexpress HH signaling proteins and neuroglial markers, we posited that these neoplasms might arise from multiple cellular sources including neuroectoderm derived precursors that are responsive to SHH signaling. In support of this premise, prior work demonstrated that selective deletion of Menin in glial cells expressing the human *GFAP* promoter or SRY-box transcription factor 10 (SOX10) was sufficient to stimulate the development of pancreatic and pituitary NETs. *GFAP-Cre; Men1^FL/FL^* (*GFAP^ΔMen1^*) mice developed NETs by 15 to 17-months of age, whereas tumors developed by 11-months of age in the *Sox10-Cre; Men1^FL/FL^* (*Sox10^ΔMen1^*) mice (*21*). We subsequently analyzed 18 to 20-month-old *GFAP^ΔMen1^* and *Sox10^ΔMen1^* mice and observed a low incidence of SI tumors arising in the antropyloric Brunner’s glands, duodenum, jejunum, and ileum. Unlike PNETs that exhibited a well-differentiated NET phenotype, SI tumors appeared to be poorly differentiated (Fig. 3A), yet these tumors were immunoreactive for the NET marker synaptophysin (SYP) (Fig 3B). Mirroring our previous observations in human PNETs and SI-NETs, tumors from the *ΔMen1* mice expressed SHH signaling pathway proteins, with PNETs showing stronger PTCH1 expression and SI tumors showing stronger SHH immunoreactivity (Fig 3B). Western blot analysis of the *ΔMen1* mouse tumors further confirmed significantly elevated expression of PTCH1, GLI1, and GLI2 proteins in PNETs compared to age-matched wild type pancreas tissues (Fig. 3, C and D). In comparison, SI tumors from the *ΔMen1* mice were significantly enriched in SHH protein expression but exhibited variable expression of GLI2 and PTCH1 proteins. Consistent with the histological findings, PNETs showed consistent and robust expression of CHGA, SYP, and PAX6. The expression of these neuroendocrine proteins was also elevated, although to a lesser extent, in SI tumors compared to wild type controls. Based on these expression patterns, we concluded that these SI tumors were bona fide SI-NETs. IHC analysis of the tumors identified absent or low nuclear Menin expression compared to adjacent and control tissues, supporting the premise that NETs develop from a Menin-deficient cell population (Fig. 3, E and F). To our knowledge, these findings represented the first report of proximal SI-NETs that develop in a genetic mouse model.

**Figure 3.**
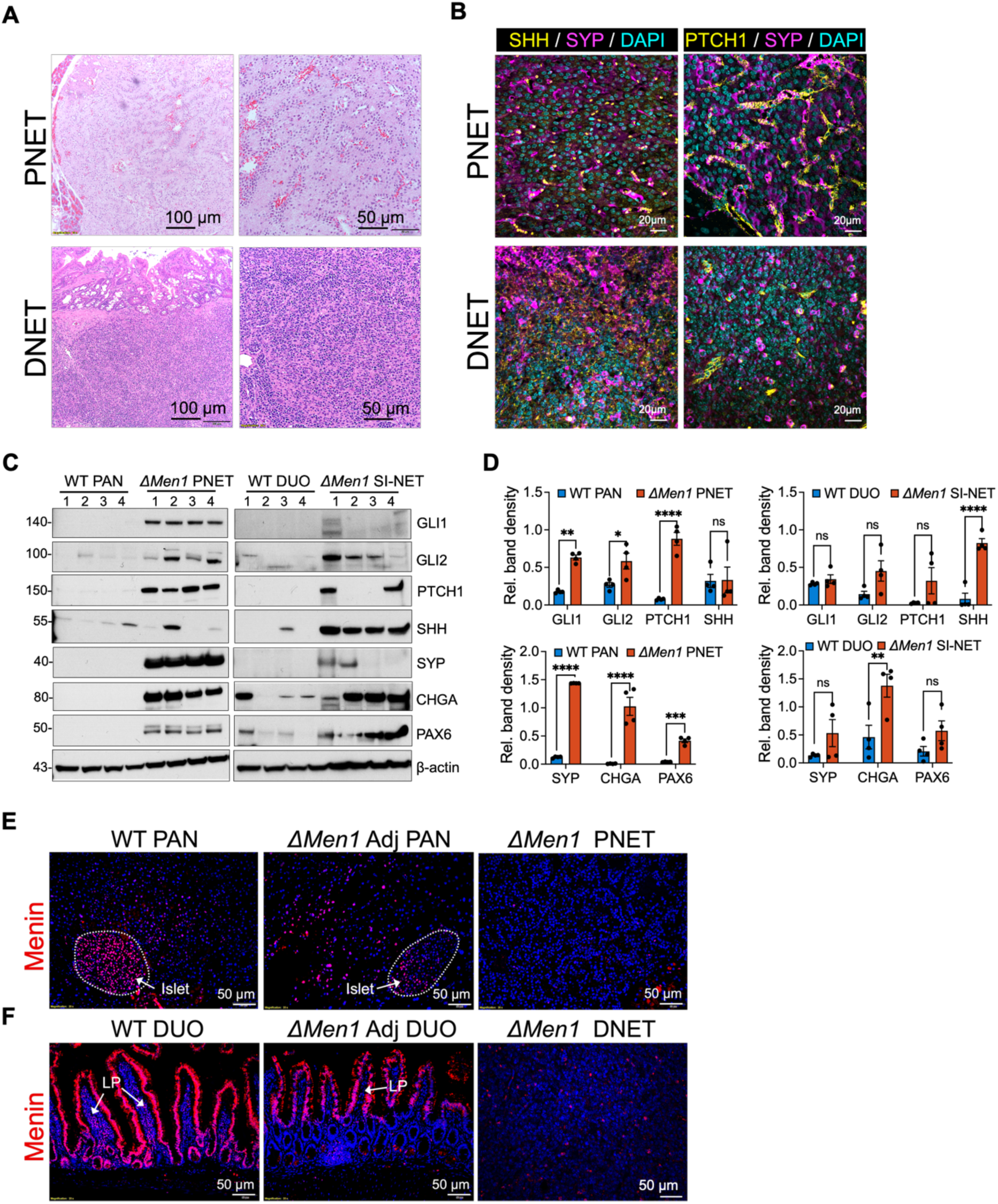
GFAP-directed *Men1* deletion stimulates pancreatic and small intestinal NETs that overexpress HH signaling proteins. (**A**) Representative hematoxylin and eosin (H&E) stained images of a pancreatic and duodenal NET (PNET in top panel and DNET in bottom panel) from 18-month-old *GFAP-Cre: Men1^FL/FL^*(*GFAP^ΔMen1^*) mice. (**B**) IF stained images of a *GFAP^ΔMen1^* PNET and DNET: Left panel shows SHH (yellow), and right panel shows PTCH1 (yellow) co-stained with SYP (magenta) and DAPI (cyan). (**C**) Western blot analysis of HH signaling proteins and neuroendocrine markers in littermate wild type pancreas (WT PAN), duodenum (WT DUO), and *GFAP^ΔMen1^* PNETs and SI-NETs. (n = 4). (**D**) Quantitation of western blot analysis in panel (C). (n = 4). * = *p* < 0.05, ** = *p* <0.01, **** = *p* < 0.0001 by Two-way ANOVA with Sidak post-test. (**E**) IF stained images showing Menin (red) expression in a *GFAP^ΔMen1^* PNET and the adjacent exocrine and endocrine (islet encircled in white dotted line) pancreas (Adj PAN). Menin expression in the endocrine islet and exocrine pancreas of a littermate control is shown for comparison. (**F**) IF stained images showing Menin (red) expression in a *GFAP^ΔMen1^* DNET and the adjacent duodenum (Adj DUO) with arrow indicating to the lamina propria (LP). Menin expression in the duodenum of a littermate control is shown for comparison. DAPI (blue).

To visualize the fate of GFAP^+^ and SOX10^+^ glial cells, we crossed the previous *GFAP^ΔMen1^* and *Sox10^ΔMen1^* mice to express the *lox-stop-lox-tdTomato* fluorescent reporter (*LSL-tdTomato*). After confirming appropriate tissue-restricted expression of the fluorescent reporter (fig. S2), we analyzed tumors from the *ΔMen1* mice and observed the presence of tdTomato^+^ signal in three mouse PNETs and in one DNET arising in the Brunner’s glands (Fig. 4, A and B). Subsequently, we generated primary tumoroids from these tdTomato^+^ *ΔMen1* tumors for further *in vitro* studies. Prior to conducting these *in vitro* experiments, we confirmed that tumoroids derived from the tdTomato^+^ *ΔMen1* tumors expressed the fluorescent reporter. Subcellular tdTomato expression in the tumoroids was consistent with nuclear SOX10 expression and localization of GFAP to the cytoplasm (Fig. 4, B and C). After confirming that the *ΔMen1* tumoroids express CHGA, we further evaluated their expression of the mitotic marker Ki-67 and HH pathway proteins. Mirroring the expression of parent mouse tumors, *ΔMen1* tumoroids demonstrated strong immunoreactivity for SHH, PTCH1, and SMO proteins (Fig. 4, E and F). Thus, similar to human GEP-NETs, *ΔMen1* PNETs and SI-NETs overexpressed HH pathway proteins. Morover, these tumors showed the involvement of tdTomato-labeled GFAP^+^ and SOX10^+^ glial cells upon loss of *Men1*.

**Figure 4.**
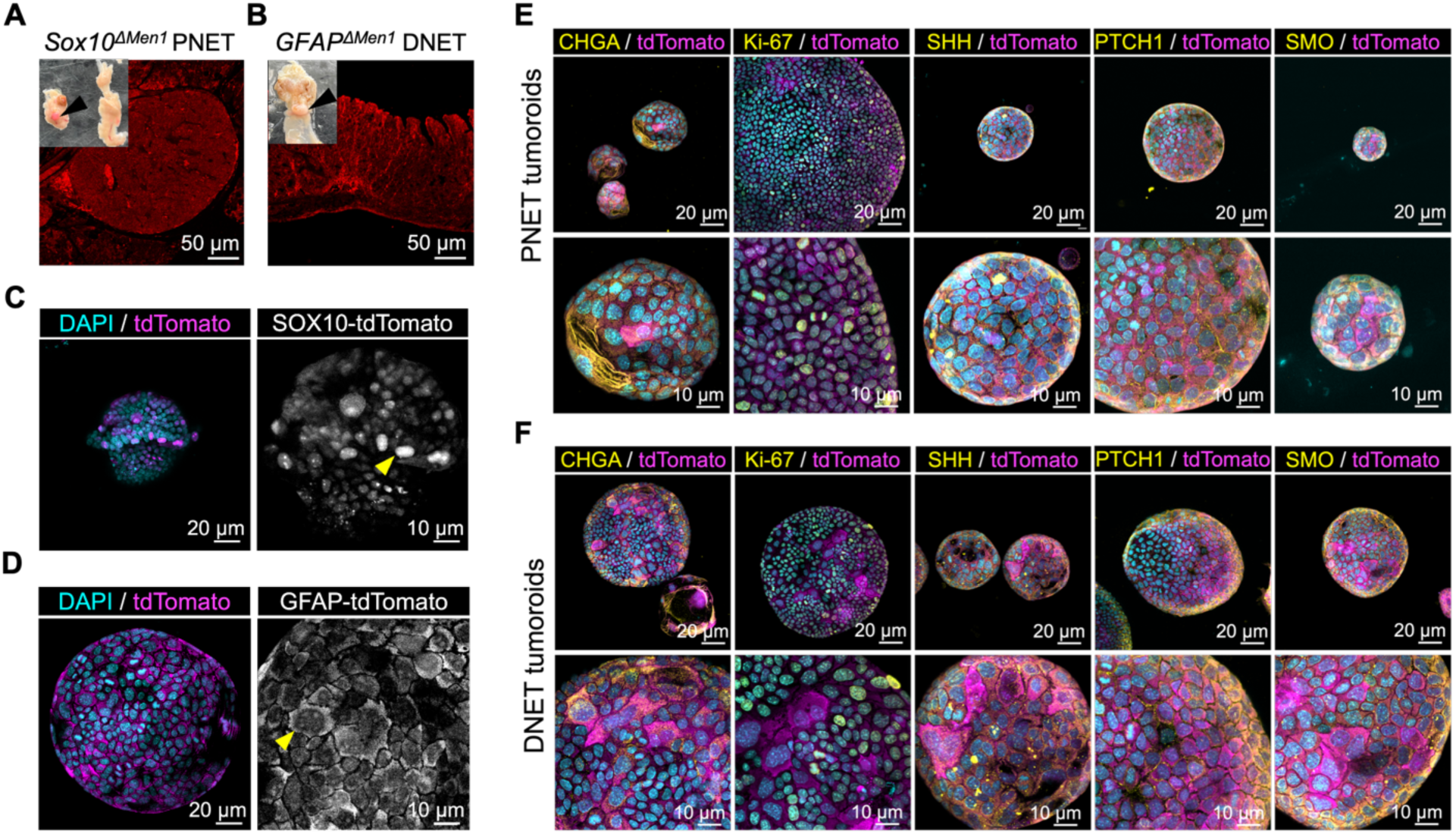
Pancreatic and small intestinal NET tumoroids from *GFAP^ΔMen1^* and *Sox10^ΔMen1^* mice recapitulate parent tumor phenotypes. (**A**) IF image showing tdTomato (red) expression in a PNET from a 17-month-old *Sox10-Cre: Men1^FL/FL^*: *LSL-tdTomato* mouse (*Sox10^ΔMen1; tdTomato^*). Inset shows the macroscopic image of the associated pancreas with two large PNETs. Arrowhead indicates to the tdTomato^+^ PNET. (**B**) IF image showing tdTomato (red) expression in a DNET from a 19-month-old *GFAP-Cre: Men1^FL/FL^*: *LSL-tdTomato* mouse (*GFAP^ΔMen1; tdTomato^*). Inset shows the macroscopic image of the associated stomach and proximal duodenum. Arrowhead indicates to the tdTomato^+^ DNET arising in the pyloric Brunner’s glands. (**C**) IF image showing tdTomato (magenta) expression in a 14-day-old tumoroid derived from a *Sox10^ΔMen1; tdTomato^* PNET. Left panel shows overlay with DAPI (cyan) and the right panel shows the tdTomato channel only. (**D**) IF image showing tdTomato (magenta) expression in a 14-day-old tumoroid derived from a *GFAP^ΔMen1; tdTomato^* DNET. Left panel shows overlay with DAPI (cyan) and the right panel shows the tdTomato channel only. (**E**) IF images of *Sox10^ΔMen1; tdTomato^* PNET tumoroids stained for CHGA, Ki-67, and HH signaling proteins (yellow) co-localized with tdTomato (magenta) and DAPI (cyan). (**F**) IF images of *GFAP^ΔMen1; tdTomato^* DNET tumoroids stained for CHGA, Ki-67, and HH signaling proteins (yellow) co-localized with tdTomato (magenta) and DAPI (cyan). Panels (*C-E*) represent maximum intensity projections of Z-stack images of the tumoroids.

### Activation of GLI1/2 regulates the growth of *ΔMen1* GEP-NET tumoroids

Unlike patient-derived tumoroids that exhibit indolent growth over subsequent passage numbers (*6,39*), the *ΔMen1* tumoroids were robust, able to be passaged and proliferated in normal organoid growth media. Thus, we leveraged these *in vivo* models to further investigate the NET cell response to SHH pathway modulation (Fig. 5A). We generated tdTomato^+^ *ΔMen1* PNET tumoroid lines from two independent mouse tumors and treated the tumoroids with HH pathway agonists, pharmacologic inhibitors of SMO, or inhibitors of GLI1/2 signaling (Fig. 5B). Consistent with the response shown by patient-derived PNET tumoroids, *ΔMen1* PNET tumoroids demonstrated increased growth upon SHH pathway activation (Fig. 5C). Compared to inhibition of the canonical pathway, pan-inhibition of SMO and GLI1/2 inhibited tumoroid growth by 50 to 90 percent (Fig. 5, D and E). These patterns remained consistent when tumoroids were treated together with respective pharmacological inhibitors and the SHH ligand (Fig. 5F). Given that patient-derived PNET tumoroids upregulated their expression of neuroendocrine transcripts in response to SHH treatment, we further tested whether *ΔMen1* PNET tumoroids undergo similar transcriptional changes in response to HH pathway modulation. Indeed, adding SHH ligand increased the expression of *Chga* in *ΔMen1* PNET tumoroids, whereas treatment with itraconazole or vismodegib suppressed expression (Fig. 5G).

**Figure 5.**
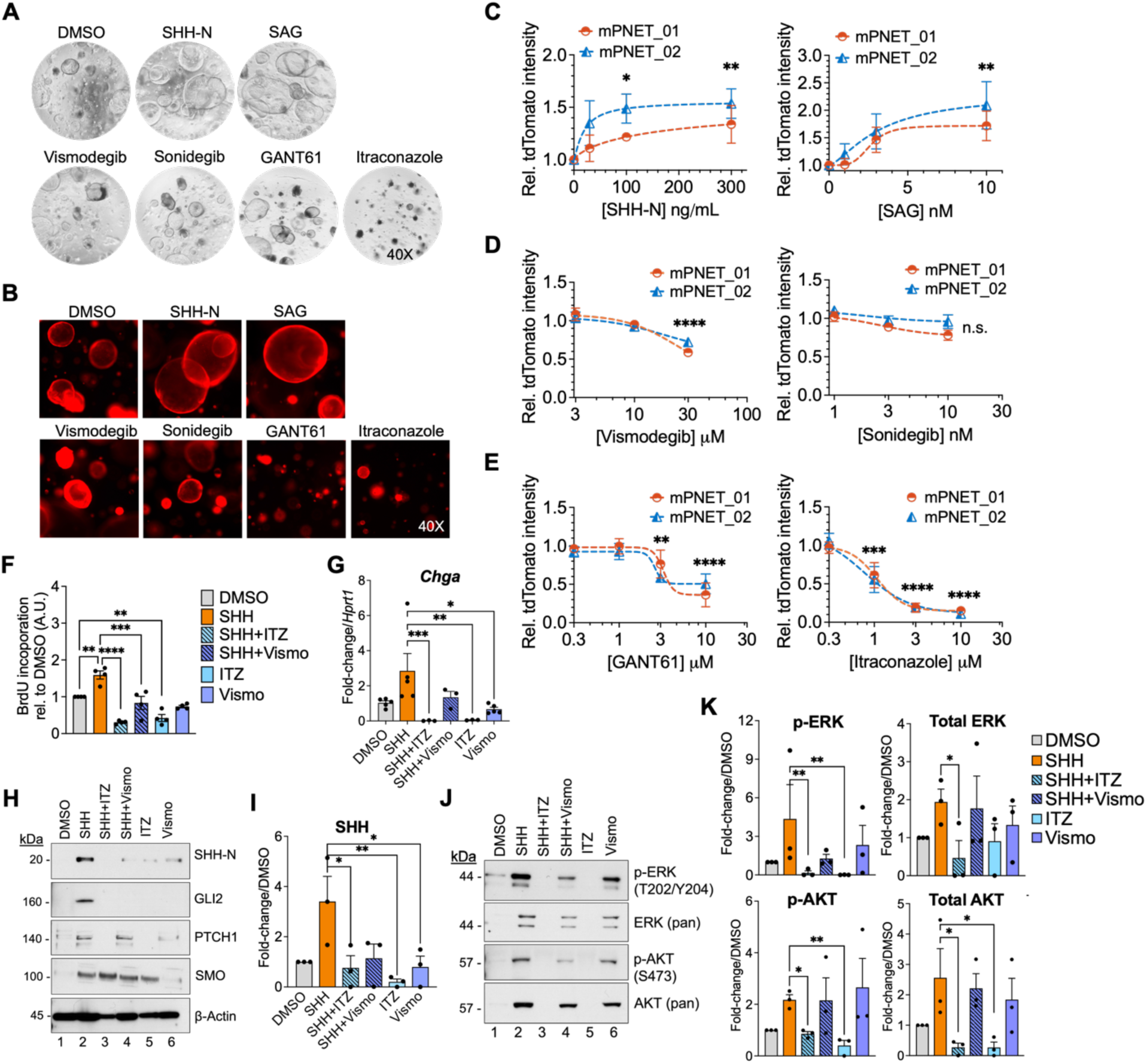
HH signaling regulates the growth of *GFAP^ΔMen1^* and *Sox10^ΔMen1^* pancreatic NET tumoroids. (**A**) Phase contrast and (**B**) fluorescence images of PNET tumoroids from *Sox10-Cre;Men1^FL/FL^*;*LSL-tdTomato* mice. Tumoroids were imaged after 72 h exposure to: the HH agonists SHH-N (100 ng/mL) and SAG (10 nM); inhibitors of the canonical HH signaling pathway vismodegib (20 µM) and sonidegib (10 nM); or inhibitors of the GLI1/2 effectors GANT61 (10 µM) and itraconazole (ITZ, 1 µM). (**C**) TdTomato fluorescence intensity was used to measure PNET tumoroid growth in the presence of HH pathway agonists or (**D**) inhibitors of the canonical and (**E**) non-canonical HH signaling pathways. Fluorescence signal is compared to DMSO vehicle control. (n = 3 replicates in two unique mouse PNET tumoroid lines). ** = *p* <0.01, *** = *p* < 0.001, **** = *p* < 0.0001 by One-way ANOVA with Dunnett post-test. (**F**) BrdU incorporation was used to measure tumoroid proliferation in *GFAP^ΔMen1^* and *Sox10^ΔMen1^* PNET tumoroids after 72 h treatment. (n = 4). ** = *p* <0.01, **** = *p* < 0.0001 by One-way ANOVA with Dunnett post-test. (**G**) Relative fold-change in *Chga* mRNA levels in PNET tumoroids following 72 h treatment. (n = 5). (**H**) Western blot analysis of HH pathway proteins in PNET tumoroids after 72 h treatment. (**I**) Quantitation of SHH protein expression normalized to beta-actin and DMSO vehicle control from the western blot analysis in panel (H). (n = 3). (**J**) Western blot analysis and (**K**) associated quantitation of phosphorylated and total ERK and AKT growth pathways in PNET tumoroids after 72 h treatment. (n = 3). * = *p* <0.05, ** = *p* <0.01, *** = *p* < 0.001 by Kruskal-Wallis test.

To confirm the downstream signaling targets of SHH pathway modulation, we measured the protein levels of SHH, GLI2, PTCH1, and SMO in *ΔMen1* PNET tumoroids following SHH pathway activation and inhibition. As anticipated, SHH treatment induced the expression of HH pathway proteins and inhibition with itraconazole, but not vismodegib, reversed the induction of these proteins (Fig. 5, H and I). Prior reports showed that SHH signaling can activate ERK and AKT through direct crosstalk with these signaling pathways (*41*). Therefore, we examined the impact of SHH pathway modulation on the expression and activation of ERK and AKT proteins in *ΔMen1* PNET tumoroids. Consistent with the previous effects on tumoroid growth, SHH treatment induced the expression of phosphorylated and total ERK and AKT, and these effects were attenuated under pan-inhibition of SMO and GLI1/2, but not when SMO was inhibited alone (Fig. 5, J and K). Collectively, these observations validated our findings in patient-derived PNET tumoroids and implicated the non-canonical SHH signaling pathway in driving tumor cell growth.

Given that human and SI-NETs overexpress SHH, we rationalized that these neoplasms might also respond to HH pathway inhibition. We repeated the previous drug studies on a DNET tumoroid line that was established from a tdTomato^+^ *ΔMen1* tumor located in the Brunner’s glands (Fig. 6, A–C). Mimicking the response by *ΔMen1* PNET tumoroids, the DNET tumoroids showed increased growth in response to SHH signaling pathway activation and inhibition of GLI1/2, but not inhibition of SMO alone, resulted in significant growth inhibition. We validated these studies using a second SI-NET tumoroid line derived from a jejunal tumor (Fig. 6D). Interestingly, SHH treatment did not lead to a significant increase in HH protein levels or the expression of ERK and AKT beyond baseline levels, suggesting that high basal expression of SHH ligand in SI-NETs contributed to autocrine growth signaling (Fig. 6, E and F). Despite exhibiting reduced sensitivity to SHH ligand, pan-inhibition of SMO and GLI1/2 was nonetheless effective in reducing tumoroid growth and downregulated the expression of ERK and AKT proteins (Fig. 6G). Finally, we compared the drug response of the *ΔMen1* DNET tumoroids to STC-1 cells derived from a mouse ileal NET (Fig. 6H). SHH and SAG treatment stimulated STC-1 cell proliferation in a dose-dependent manner and inhibition of both canonical and non-canonical HH pathways resulted in the opposite effect (Fig. 6, I–K). These results indicated that STC-1 cells are characterized by increased sensitivity to modulation of the canonical HH signaling pathway, whereas primary SI-NET tumoroids more closely mimicked the drug response shown by patient-derived PNET tumoroids.

**Figure 6.**
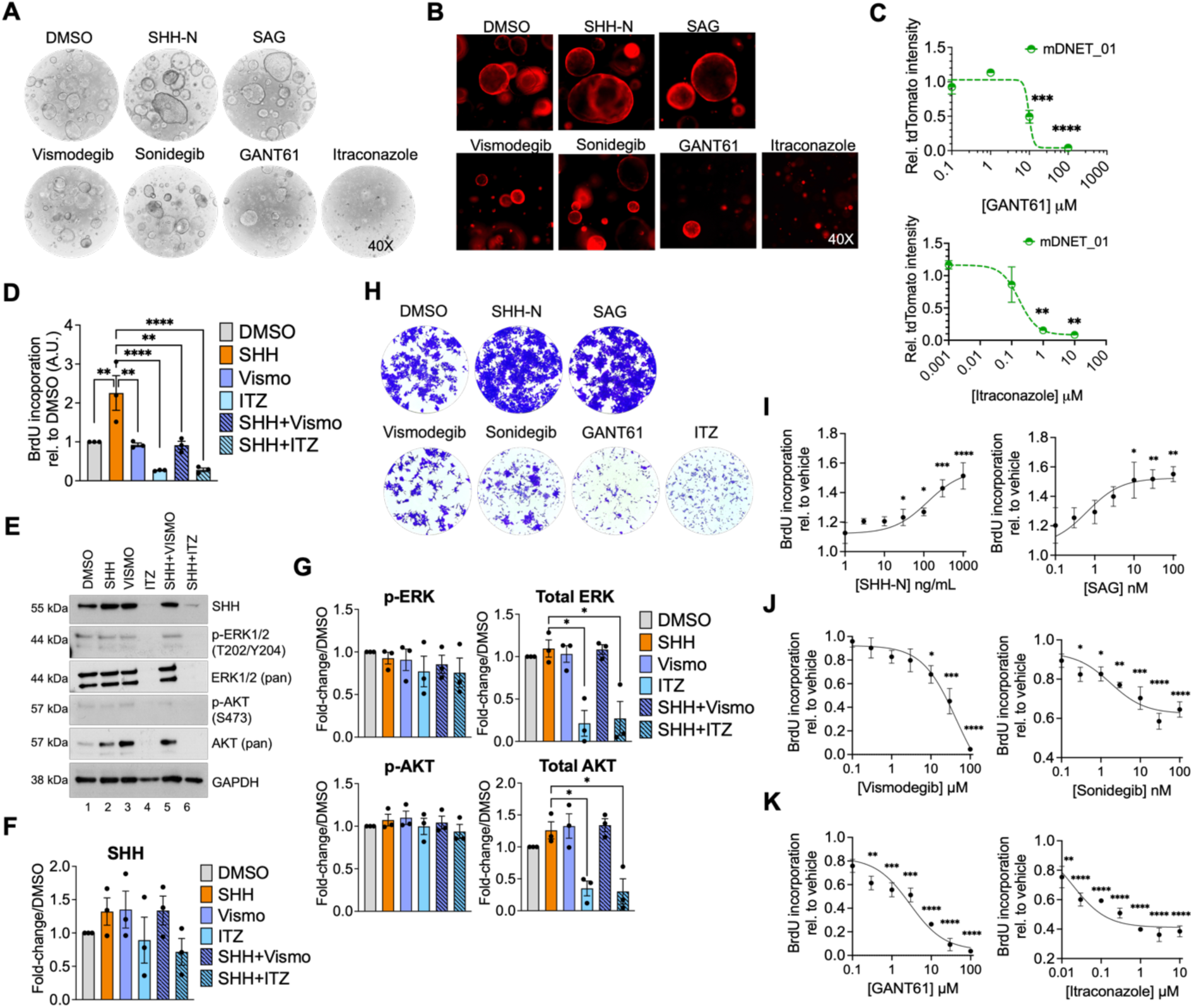
Small intestinal NETs are sensitive to HH pathway activation and inhibition. (**A**) Phase contrast and (**B**) fluorescence images of SI-NET tumoroids from a *GFAP-Cre;Men1^FL/FL^*;*LSL-tdTomato* mouse. Tumoroids were imaged after 72 h exposure to: the HH agonists SHH-N (100 ng/mL) and SAG (10 nM); inhibitors of the canonical HH signaling pathway vismodegib (20 µM) and sonidegib (10 nM); or inhibitors of the GLI1/2 effectors GANT61 (10 µM) and itraconazole (ITZ, 1 µM). (**C**) TdTomato fluorescence intensity was used to measure SI-NET tumoroid growth in the presence of HH pathway inhibitors. Fluorescence signal is compared to DMSO vehicle control. (n = 3 replicates using one mouse SI-NET tumoroid line). ** = *p* <0.01, *** = *p* < 0.001, **** = *p* < 0.0001 by One-way ANOVA with Dunnett post-test. (**D**) Relative BrdU incorporation by *GFAP^ΔMen1^* and *Sox10^ΔMen1^* PNET tumoroids after 72 h treatment. (n = 4). ** = *p* <0.01, *** = *p* < 0.001, **** = *p* < 0.0001 by One-way ANOVA with Dunnett post-test. (**E**) Western blot analysis of SHH, ERK, and AKT growth pathways in SI-NET tumoroids after 72 h treatment. (**F**) Western blot quantitation of SHH and (**G**) phosphorylated and total ERK and AKT proteins normalized to GAPDH and DMSO vehicle control. (n = 3). * = *p* <0.05 by Kruskal-Wallis test. (**H**) Crystal violet staining of mouse STC-1 SI-NET cells after 48 h treatment with agonists and inhibitors of the HH signaling pathway. (**I**) BrdU incorporation in STC-1 cells after 48 h treatment with HH agonists SHH-N and SAG, (**J**) SMO inhibitors vismodegib and sonidegib, and (**K**) inhibitors of GLI1/2 signaling. * = *p* < 0.05, ** = *p* <0.01, *** = *p* < 0.001, **** = *p* < 0.0001 by One-way ANOVA with Dunnett post-test.

### Reciprocal signaling by Menin and SHH regulate enteric glial cell identity

Although our previous observations in the tdTomato^+^ *ΔMen1* mice indicated that PNETs and SI-NETs can develop from GFAP^+^ and SOX10^+^ enteric glial cells (EGCs), the mechanisms leading up to these events remained undefined. To determine how EGCs respond to *Men1* deletion, we generated primary mouse EGC cultures from the proximal duodenum of *Sox10-CreER^T2^-LSL-tdTomato* mice. TdTomato fluorescence was induced by treating the cells with 4-hydroxytamoxifen (4-OHT) and the resulting tdTomato^+^ EGCs were purified by fluorescence activated cell sorting (FACS) to yield a pure SOX10^+^ cell population (Fig. 7, A–C). We next knocked down the expression of *Men1* in EGC subcultures using small interfering RNAs and evaluated the expression of HH signaling pathway genes (Fig. 7D). *Men1* silencing resulted in significant upregulation of *Shh* and *Gli1* transcripts, and this was consistent with increased protein expression (Fig. 7, E–G). We further evaluated whether *Men1* knockdown led to reprogramming of EGCs by altering the expression of glial and neuroendocrine lineage transcripts. Indeed, we observed a downregulation in several glial lineage-restricted genes (*e.g., Gfap, S100b, Fabp7,* and *Lpar1*) and upregulated expression of neuroendocrine and neural progenitor transcripts that are also enriched in GEP-NETs (*e.g, Chga, Pax6,* and *Ascl1*) (Fig. 7, H and I). To test whether these events are mediated by upregulated SHH signaling in response to loss of *Men1*, we treated EGCs with GANT61 or vismodegib and measured changes in the expression of HH and neuroendocrine genes. As predicted, inhibition of GLI1/2 with GANT61 reversed the induction of HH transcripts and this coincided with downregulated expression of *Chga* and *Pax6* (Fig 7, J and K). In comparison, SMO inhibition with vismodegib did not significantly alter the expression of neuroendocrine transcripts in *Men1*-deleted cells. Given that Menin was reported to antagonize HH signaling (*23,24*), we evaluated whether GLI1/2 inhibition resulted in reciprocal feedback on Menin expression in EGCs. Interestingly, GANT61 treatment rescued nuclear Menin expression in EGCs within three days of *Men1* silencing, suggesting that Menin and the GLI1/2 effectors participate in reciprocal regulation (Fig. 7L). In summary, these studies demonstrated that EGCs reprogram from a glial-restricted lineage and acquire a neuroendocrine phenotype in response to *Men1*-mediated HH pathway activation.

**Figure 7.**
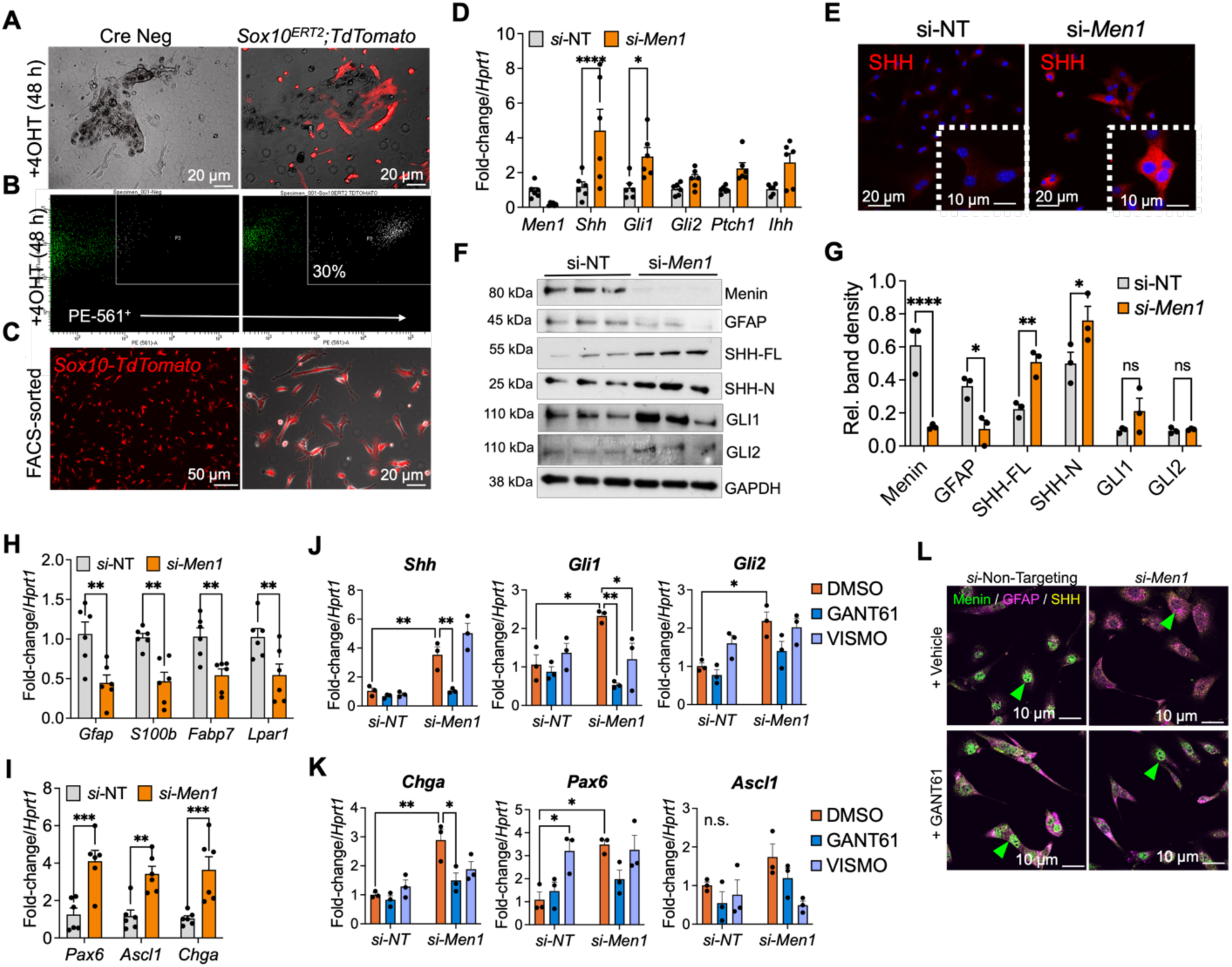
Loss of *Men1* in enteric glial cells stimulates GLI1/2-dependent transcriptional reprogramming. (**A**) Combined fluorescence and phase contrast images of 5-day-old primary enteric glial cell (EGC) cultures from *Sox10-CreER^T2^*;*LSL-tdTomato* mice and *CreER^T2^* negative controls. Top panel shows TdTomato+ EGCs after 48 h exposure to 4-hydroxytamoxifen 4-OHT (2 µM). (**B**) TdTomato+ EGCs were sorted by FACS to enrich for a pure SOX10+ cell population. (**C**) Combined fluorescence and phase contrast images of FACS-enriched SOX10-tdTomato+ EGCs. (**D**) Fluctuations in HH pathway mRNA levels were evaluated in SOX10-tdTomato+ EGCs 72 h following siRNA-mediated *Men1* silencing. siRNA treatment consisted of four pooled siRNAs targeting the *Men1* gene (*si-Men1*, 25 µM) or non-targeting (si-NT, 25 µM) controls. (n = 5). (**E**) Immunofluorescence images of SHH expression in si-NT and *si-Men1* treated EGCs (SHH = red pseudo-color, DAPI = blue). Inset shows higher power image. (**F**) Western blot analysis of si-NT and *si-Men1* EGCs after 72 h treatment. SHH-FL = 55 kDa full length peptide; SHH-N = 22 kDa N-terminal peptide. (n = 3). (**G**) Quantitation of protein expression in panel (F) normalized to GAPDH loading control. (n = 3). (**H**) Relative fold-change in glial lineage transcripts and (**I**) neuroendocrine and neural progenitor transcripts in si-NT and *si-Men1* treated EGCs. (n = 6). (**J**) Immunofluorescence images of si-NT and *si-Men1* EGCs after 96 h siRNA knockdown and 72 h treatment with vehicle or GANT61 (10 µM). Menin = green, GFAP = magenta, SHH = yellow. (**K**) qPCR analysis of *Men1* expression in si-NT and *si-Men1* EGCs after 72 h treatment with GANT61 (10 µM) or vismodegib (VISMO 20 µM). (n = 3). (**L**) Relative fold-change in HH pathway genes and (**M**) neuroendocrine and neural progenitor transcripts in si-NT and *si-Men1* EGCs after treatment with GANT61 or VISMO (n = 3). For all plots, * = *p* < 0.05, ** = *p* <0.01, *** = *p* < 0.001, **** = *p* < 0.0001 by Two-way ANOVA with Sidak post-test.

### SHH signaling in GFAP^+^ glia is required for GEP-NET development

Our studies using primary EGCs indicated that pharmacologic inhibition of downstream SHH signaling effectors could reverse the glial-to-neuroendocrine transition. The extent to which these events occur in vivo and their impact on GEP-NET development remained unexplored. Thus, we sought to translate our in vitro observations by generating *GFAP^ΔMen1^* mice that harbored *Kif3a*-deficient EGCs showing impaired SHH signaling (*GFAP^ΔMen1; ΔKif3a^*). *Kif3a* encodes a structural protein for primary cilia, a required component of SHH signal transduction present on many stromal cell types including those derived from the neural crest (*42–45*). We evaluated mice aged 18-to 20-months for the presence of NETs and observed a remarkable reduction in the number of PNETs and SI-NETs in the *GFAP^ΔMen1; ΔKif3a^* mice compared to mice with an intact *Kif3a* locus (Fig. 8, A–D). Consistent with a reduction in tumors, the *GFAP^ΔMen1; ΔKif3a^* mice exhibited significantly lower expression of circulating pancreatic and gastrointestinal hormones (Fig. 8, F–H). Intriguingly, the incidence of pituitary NETs and the expression of prolactin, the dominant hormone expressed by these tumors (*21*), were not affected by *Kif3a* deletion (Fig. 8, I and J). This suggested that unlike NETs that developed in the GI tract, the development of pituitary NETs did not depend on activated SHH signaling in GFAP^+^ glia.

**Figure 8.**
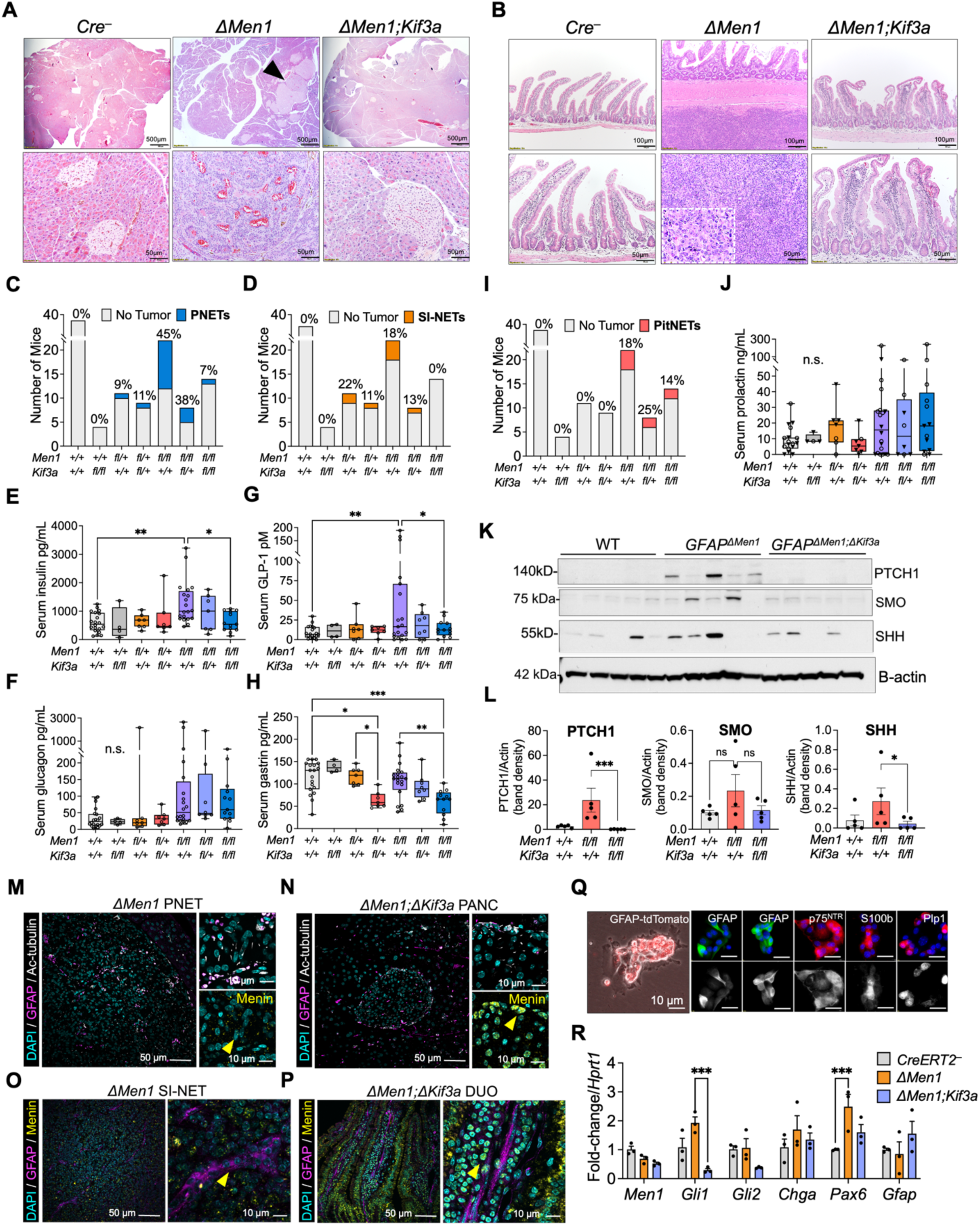
Genetic inhibition of SHH signaling in *GFAP^ΔMen1^* mice abolishes the development of pancreatic and small intestinal NETs. (**A**) Representative H&E images of pancreas from 19-month-old littermate wild type controls (*Cre^−^*), *GFAP^ΔMen1^* mice, and *GFAP^ΔMen1;Kif3a^* mice. Arrowhead indicates to PNET. Higher magnification images are shown in the bottom panel. (**B**) Representative H&E images of duodenum from 19-month-old littermate wild type controls (*Cre^−^*), *GFAP^ΔMen1^* mice, and *GFAP^ΔMen1;Kif3a^* mice. Arrowhead indicates to PNET. Higher magnification images are shown in the bottom panel and inset. (**C**) Incidence of PNETs and (**D**) incidence of SI-NETs in 17 to 20-month-old littermate wild type controls, compared to *GFAP^ΔMen1^* mice and *GFAP^ΔMen1;Kif3a^* mice with heterozygous and homozygous allelic deletion. (**E**) Fasting serum insulin, (**F**) glucagon, (**G**) GLP-1, and (**H**) gastrin levels in 17 to 20-month-old littermate wild type controls, *GFAP^ΔMen1^* mice, and *GFAP^ΔMen1;Kif3a^* mice. (n = 4–20 mice per genotype). * = *p* < 0.05, ** = *p* <0.01, *** = *p* < 0.001, **** = *p* < 0.0001 by One-way ANOVA with Dunnett post-test. (**I**) Incidence of pituitary NETs (PitNET) in 17 to 20-month-old littermate wild type controls, compared to *GFAP^ΔMen1^* mice and *GFAP^ΔMen1;Kif3a^* mice. (**J**) Fasting serum prolactin levels in 17 to 20-month-old mice stratified by sex: males (inverted solid black triangle) and females (open circles). (**K**) Western blot analysis of HH signaling proteins in *GFAP^ΔMen1^* PNETs compared to pancreas from *GFAP^ΔMen1;Kif3a^* mice and littermate WT controls. (n = 5 mice per genotype). (**L**) Quantitation of western blots in panel (K). (n = 5 mice per genotype). * = *p* < 0.05, *** = *p* < 0.001 by Kruskal-Wallis test. (**M**) IF images of a *GFAP^ΔMen1^* PNET and (**N**) *GFAP^ΔMen1;Kif3a^* pancreas stained for GFAP (magenta), the ciliary marker acetylated tubulin (Ac-tubulin), and DAPI (cyan). Higher magnification image is shown in the top right panel. Bottom right panel shows Menin (yellow) co-localized with DAPI (cyan). (**O**) IF images of a *GFAP^ΔMen1^* DNET and (**P**) *GFAP^ΔMen1;Kif3a^* duodenum stained for GFAP (magenta), the ciliary marker acetylated tubulin (Ac-tubulin), and DAPI (cyan). Higher magnification image is shown in the top right panel. Bottom right panel shows Menin (yellow) co-localized with DAPI (cyan). (**Q**) Overlay of an IF and phase-contrast image showing 4-day-old enteric glial cell (EGC) cultures from *GFAP^tdTomato^* mice (left panel). Right panels show EGCs stained for glial cell markers GFAP (green), p75NTR (red), S100B (red), and PLP1 (red) co-localized to DAPI (blue). (**R**) Four-day-old primary EGCs from tamoxifen-inducible *GFAP-CreER^T2;ΔMen1;Kif3a^* mice were exposed to 4-hydroxy-tamoxifen (4-OHT) for 48 hours and analyzed for changes in relative mRNA abundance. (n = 3). *** = *p* < 0.001, by Two-way ANOVA with Sidak post-test.

We further confirmed downregulated expression of HH proteins in tissues from the *GFAP^ΔMen1; ΔKif3a^* mice (Fig. 8, K and L). These results were consistent with the presence of shortened primary cilia surrounding the endocrine islets of these mice compared to the cilia expressed on *GFAP^ΔMen1^* PNETs (Fig. 8, M and N) (*21*). Further IHC analysis of these tissues showed that nuclear Menin expression was lost in *GFAP^ΔMen1^* PNETs, whereas the endocrine islets of *GFAP^ΔMen1; ΔKif3a^* mice showed positive nuclear Menin expression. We next analyzed SI-NETs and the proximal intestine from the respective genetic mice and observed a similar rescuing of nuclear Menin expression in the lamina propria of the intestinal mucosa (Fig. 8, O and P). Finally, to confirm the specificity of these events, we generated primary EGCs from *GFAP-CreER^T2ΔMen1; ΔKif3a^* mice and validated their expression of glial proteins (Fig. 8Q). Following induction with 4-OHT, we observed a reduction in *Gli1* and *Gli2* transcript levels that coincided with decreased *Pax6* expression in EGCs from the *GFAP-CreER^T2ΔMen1; ΔKif3a^* mice (Fig. 8R). Collectively, these studies demonstrated that activation of HH signaling in GFAP^+^ EGCs is a required step in *ΔMen1* GEP-NET formation and underscores a glial cell of origin for these tumors.

## DISCUSSION

Augmented SHH signaling, either through canonical SMO signal transduction or non-canonical activation of the transcriptional effectors GLI1/2, promotes cellular growth in pancreatic adenocarcinoma (PDAC), basal cell carcinoma (BCC), medulloblastoma, and gastroenteropancreatic neuroendocrine tumors (GEP-NETs) (*23,33,37–39*). Targeting HH clinically has proven more nuanced, with HH inhibitors prolonging overall survival in clinical trials of BCC and medulloblastoma and accelerating disease progression in certain cases of PDAC (*37,38,46–48*). This paradoxical response to HH inhibition is attributed to fundamental differences in how these tissues respond to HH pathway activation and whether these signals function through cell autonomous, non-cell autonomous, canonical, and non-canonical signaling pathways. In cancers with neuroectoderm origin, such as BCC and medulloblastoma, tumor secreted SHH acts in an autocrine fashion to promote cellular transformation, whereas endoderm derived cancers including PDAC, rely on paracrine SHH signals that can stimulate or restrain cell growth (*49,50*). Thus, identifying the cellular sources of SHH signals and how they drive malignant progression in tissues with ectodermal and endodermal origins are essential to designing effective clinical therapies and underscores the importance of this study.

Our findings confirm a pro-tumorigenic role for activated HH signaling in GEP-NET development and point to the potential promise of targeting aberrant SHH activation in the clinic. Current FDA-approved compounds that target this pathway are restricted to SMO inhibitors, which were shown in our study to have limited therapeutic effect compared to direct inhibitors of the GLI1/2 transcription factors. Hence, our findings warrant a reassessment of pharmacologic inhibitors of the non-canonical HH signaling pathway in targeting these tumors. We further elucidate on a mechanism for HH activation in GEP-NETs and establish the role of SHH in stimulating the transformation of EGCs, a stromal cell population that demonstrates transcriptional plasticity and represents an alternative etiology for these cancers. Our results suggest that HH overactivation in GEP-NETs mimics the pattern of SHH signaling in neuroectoderm derived cancers, and inhibition of this pathway poses therapeutic benefit by restricting the reprogramming of glial cells into hormone-producing NETs.

Recent multiome sequencing demonstrated that EGCs maintain lineage plasticity and can give rise to a neurogenic cell lineage, yet the signaling cues that regulate these differentiation events remain unknown. GFAP^+^ glial cells of the central nervous system were previously shown to reprogram into neural progenitor cells and functional neurons in the presence of activated SHH signaling (*29,51*). Further, SHH pathway activation upregulated the expression of neural progenitor cell (NPC) transcription factors (*e.g.,* PAX6, ASCL1, and SOX2) that support glial-to-neural reprogramming and are essential for neuroendocrine cell specification (*52–54*). Although these events are well defined in CNS glia, pathological activation of HH signaling in EGCs that promotes a similar NPC program has not been tested since they are not considered to be a source of cancer. Our findings address this gap in knowledge by showing that EGCs can reprogram toward a neuroendocrine lineage in response to *Men1*-mediated activation of the HH signaling pathway.

Finally, we point to several limitations in the present study that limit a broader interpretation of these findings. First, genetic deletion in the transgenic mice was accomplished using a constitutive *Cre* driver, whereas the use of a tamoxifen-responsive *CreER^T2^* system would enable a true lineage trace of tdTomato^+^ EGCs after *Men1* deletion. The use of an inducible *CreER^T2^* system would allow for precise temporal control of when *Men1* is deleted and could therefore answer whether differentiated GFAP^+^ and SOX10^+^ cells in juvenile and adult mice can undergo similar transcriptional reprogramming. Additional application of single cell profiling methods would enable a better dissection of the reprogrammed cell populations and support the identification of potential biomarkers. Second, our use of the *Kif3a*-null mice demonstrated that SHH signal transduction via primary cilia is a required step in *Men1*-driven tumor formation, however this model did not provide context into the role of canonical and non-canonical HH signaling pathways. This knowledge is relevant given that patient and mouse tumoroids showed disparate responses to pharmacologic inhibitors of SMO and GLI1/2.

Our in vitro studies using mouse EGCs were limited to cultures derived from the proximal SI, since these cells are better characterized and can be isolated using established protocols (*55,56*). EGCs in the pancreas are more enigmatic but are known to comprise Schwann cells that both encapsulate and co-localize with the endocrine islet (*57,58*). Last, our work using patient-derived tumoroids were restricted to PNETs since these tumors were more readily available. Our attempt to generate tumoroid cultures from a lone DNET was unsuccessful, mirroring the high failure rate that others have reported for SI-NETs (*6,40*). Future work must address the lack of preclinical humanized SI-NET models to better translate these findings to the clinic. For instance, the application of patient-derived xenograft models for SI-NETs may represent a promising approach to propagate slow-growing tumors and test the delivery of GLI1/2 inhibitors in future preclinical studies.

## MATERIALS AND METHODS

### Study Design

The primary objective of this study was to determine how overactivation of the SHH signaling pathway contributes to the etiology of GEP-NETs. We used archived specimens from patients with PNETs and DNETs to confirm the expression of SHH signaling proteins in tumors. We tested the therapeutic impact of HH inhibitors using patient-derived NET tumoroids and tumoroid cultures derived from mouse GEP-NETs. We used primary mouse EGCs and genetic mice to determine the glial cell response to *Men1*-mediated SHH activation. All experiments were independently replicated between three to five times. Replicates and sample sizes of patients and mice are defined in figures or figure legends. The experimental methods were performed as described in Supplementary Materials and Methods.

### Patient Samples

Formalin-fixed and paraffin-embedded (FFPE) sections of surgically resected human tumors were provided by the University of Arizona Tissue Acquisition Shared Resource (TACMASR), University of Michigan Endocrine Oncology Repository, and Biorepository and Pathology Core at Icahn School of Medicine at Mount Sinai, under Human Institutional Review Board (IRB) approvals HUM00115310 and STUDY-12-00145. Fresh surgically resected tumor tissues were provided by the University of Arizona TACMASR. Informed patient consent was provided prior to tissue collection.

### Animal Studies

Mouse experiments were performed in compliance with the University of Arizona Institutional Animal Care and Use Committee guidelines. Group sample sizes were chosen using the records of variance in past analyses. All of the experiments included similar ratios of male and female mice and included *Cre* negative littermate mice as controls for the respective genotypes. Additional details on mouse strains and housing conditions are described in the Supplementary Materials.

### Statistical Analysis

Statistical analyses were performed using GraphPad Prism 10 software. Two-tailed Student’s t test and one-way or two-way analysis of variance (ANOVA) with pertinent post-tests were applied to normally distributed datasets. Datasets with a non-normal sample distribution were analyzed using the non-parametric Kruskal-Wallis test. Error bars indicate means ± SEM. All *P* values are defined as \**P* < 0.05, ** *P* < 0.01, *** *P* < 0.001, and **** *P* < 0.0001.

## Acknowledgments

We are grateful to Dr. Tobias Else, Dr. Michelle Kim, the Biorepository and Pathology Core laboratory team at Icahn School of Medicine at Mount Sinai, and biorepository participants for their contributions to this research.

## Author contributions

Conceptualization: SD, JLM

Methodology: SD, JLM

Investigation: SD, ABT, AZ, RAS, JLM

Visualization: SD, JLM

Funding acquisition: SD, JLM

Project administration: JLM

Supervision: SD, JLM

Writing – original draft: SD, JLM

Writing – review & editing: SD, AZ, JLM

All authors reviewed and approved the manuscript.

## Data and materials availability

Individual level data associated with the present study will be made available upon reasonable request to the corresponding author. Data associated with The Cancer Genome Atlas (TCGA) analyses are publicly available through the National Cancer Institute’s Genome Data Commons Portal.

