## Supplementary Materials for "Hedgehog signaling drives glial cell plasticity and oncogenic reprogramming in gastroenteropancreatic neuroendocrine neoplasms"

Duan *et al.*

**The PDF file includes:**
Materials and Methods
Figs. S1 and S2
Tables S1 and S2

### Materials and Methods

#### Animals

Mice were housed in individually ventilated cages with food and water *ad lib.* For serum hormone analysis, the mice were fasted for 16 h with water *ad lib* prior to blood collection. The mouse strains used in the study were obtained from Jackson Laboratories and are as follows: *FVB-Tg(GFAP-cre)25Mes/J*, Strain # 004600, RRID: IMSR_JAX:004600; *129S(FVB)-Men1$^{tm1.2Ctre}$/J*, Strain # 005109, RRID: IMSR_JAX:005109; *B6.Cg-Gt(ROSA)26Sor$^{tm14(CAG-tdTomato)Hze}$/J*, Strain # 007914, RRID: IMSR_JAX:007914; *B6;CBA-Tg(Sox10-cre)1Wdr/J*, Strain # 025807, RRID: IMSR_JAX:025807; *B6.Cg-Tg(GFAP-cre/ERT2)505Fmv/J*, Strain # 012849, RRID: IMSR_JAX:012849; *CBA;B6-Tg(Sox10-icre/ERT2)388Wdr/J*, Strain # 027651, RRID: IMSR_JAX:027651. Transgenic *Cre* and *CreER$^{T2}$* expressing mice were bred onto a *Men1$^{FL/FL}$* background to conditionally delete *Men1* in GFAP- or SOX10-expressing cells. A subset of mice carried a somatostatin-null genetic background (*Sst$^{-/-}$*) and any comparisons between genotypes accounted for the presence of the null allele, as reported on previously *(20)*. *GFAP$^{\Delta Men1}$* mice were bred onto a *Kif3a$^{FL/FL}$* background. The respective genetic mice were bred to express the *lox-Stop-lox-tdTomato* sequence to selectively express the tdTomato fluorescent reporter in GFAP$^{+}$ or SOX10$^{+}$ cells. All strains were analyzed at 18 to 22 months of age. All histological characterization and downstream comparisons were made using littermate controls that genotyped negative for *Cre* recombinase expression and included mice with and without the relevant floxed alleles.

#### The Cancer Genome Atlas (TCGA) Patient Cohort Analysis

TCGA data were accessed through the National Cancer Institute's Genome Data Browser. Patient cohorts were designed using the GDC Cohort Builder and filtered based on the primary cancer diagnosis. The TCGA-neuroendocrine cohort consisted of 841 cases of which survival data was only available for 17 cases. The primary neuroendocrine diagnoses were as follows: atypical carcinoid tumor (n = 28), carcinoid tumor, nos (n = 18), extra-adrenal paraganglioma, malignant (n = 6), extra-adrenal paraganglioma, nos (n = 18), large cell neuroendocrine carcinoma (n = 86), medullary carcinoma, nos (n = 30), merkel cell carcinoma (n = 50), neuroendocrine carcinoma, nos (n = 427), paraganglioma, nos (n = 22), pheochromocytoma, malignant (n = 41), pheochromocytoma, nos (n = 126), and pituitary adenoma, nos (n = 7). Cases identified as having glial cell origins included the following primary diagnoses: astrocytoma, anaplastic (n = 131), astrocytoma, nos (n = 256), ependymoma, nos (n = 17), Glioblastoma (n = 1,400), glioblastoma multiforme (n = 18), glioma, malignant (n = 60), oligodendroglioma, anaplastic (n = 79), and oligodendroglioma, nos (n = 192). Copy number variations were analyzed by filtering for the molecular biomarkers indicated in each table or heatmap. Tables and heatmaps were generated in the GDC data portal and edited for clarity using the Inkscape open access graphic and design editor.

**Immunohistochemical/Immunofluorescence (IHC/IF) Staining**

Five-micron formalin-fixed and paraffin-embedded (FFPE) tissue sections were baked at 65°C for 60 min and then deparaffinized in three xylene washes for a total of 15 min. Tissues were rehydrated in 100%, 90%, and 70% ethanol washes, and then rinsed with Tris Buffered Saline (TBS) for 5 min. Slides were subjected to heat-mediated antigen retrieval in Tris-EDTA buffer (pH 9.0, Cat #ab93684, Abcam) for 30 min at 95°C. Slides were allowed to cool to 24°C for 20 min and then rinsed for 1 min in running distilled water followed by three 1 min washes in TBS. Slides were blocked in 10% donkey serum prepared in 1% bovine serum albumin (BSA), 0.2% Triton X-100, and 0.05% Tween-20 (TBST) for 1 h at 24°C. For mouse tissues that were subject to a primary antibody that was raised in a mouse host, sections were also blocked for 1 h using the Mouse on Mouse (M.O.M) staining kit (Vector Laboratories). Sections were then incubated in the respective primary antibodies overnight at 4°C. Slides were washed three times by submerging in TBST and incubating with gentle rocking for 5 min. The slides were next incubated in Alexa Fluor-conjugated secondary antibodies diluted 1:400 in TBST with 1% BSA (Thermo Fisher Scientific, Waltham, MA). Slides were incubated for 10 min in 4', 6-diamidino-2-phenylindole (DAPI) diluted 1:5000 in TBS, then mounted using Fluormount-G (Thermo Fisher Scientific).

For menin staining that required an additional tyramide signal amplification (TSA) step, slides were first incubated in BLOXALL reagent for 10 min to quench endogenous peroxidase activity (Vector Laboratories). The slides were washed in TBS and then blocked and processed as described above. After the overnight incubation, the slides were incubated in anti-Mouse IgG secondary antibody conjugated to Horse Radish Peroxidase (ImmPACT DAB, Vector Laboratories, Newark, CA) for 30 min at 24°C. TSA was performed according to the manufacturer's instructions using a 1:50 dilution of the fluorophore following reconstitution in dimethyl sulfoxide (DMSO) (Cat #NEL701A001KT, PerkinElmer, Waltham, MA). Slides were incubated in the TSA reagent for 5-7 min, washed, and then counterstained, if additional primary antibodies were used. IHC/IF stained slides were imaged using the Olympus BX53F epifluorescence microscope (Center Valley, PA) or with the Nikon AX R Laser-Scanning Confocal Microscope (Tokyo, Japan). Image acquisition settings were kept consistent when any direct comparisons were made. Post hoc image analyses were performed using open-source Image J (FIJI) software, were applied globally (*i.e.,* to the entire image file), and were restricted to channel level adjustments and assignment of pseudo colors.

**Immunocytochemistry (ICC/IF) Staining**

Primary tumoroids embedded in Matrigel and seeded in 24-well black glass bottom plates were washed twice with pre-warmed Dulbecco's PBS (DPBS). The tumoroids were fixed in pre-warmed 4% paraformaldehyde (PFA) for 40 min at 24°C. Organoids were permeabilized in 0.5% Triton X-100 for 20 min at 24°C. The tumoroids were washed twice with TBS and then blocked in 10% donkey serum and 1% BSA for 1 h at 24°C. The tumoroids were incubated in respective primary antibodies for 16 h at 4°C. The tumoroids were washed four times with TBST and incubated in Alexa Fluor-conjugated secondary antibodies, followed by DAPI, as described above. Tumoroids were stored in TBS prior to and during imaging using the Nikon AX R Laser-Scanning Confocal Microscope (Tokyo, Japan). Z-stack images were acquired at 100X and 200X magnification and combined into a maximum intensity projection (MIP) image using the Nikon NIS-Elements software. Pseudocoloring and adjustments to brightness levels were applied globally using ImageJ (FIJI) software.

Primary EGCs were seeded in 24-well glass bottom plates or in 6-well plates containing a glass cover slip. In both scenarios, poly-L-lysine was used to coat the plate or coverslip. Cells were stained according to the steps described above, with the following changes: cells were incubated in 4% PFA for only 15 min and in 0.5% Triton X-100 for 5 min. For crystal violet staining of the BON-1 and STC-1 cells, the cells were fixed in the same manner and washed in PBS. Next, the cells were incubated in 0.1% crystal violet dye for 20 min at 24°C. Cells were washed with distilled water and allowed to dry prior to imaging on the ECHO Revolve inverted microscope (ECHO, San Diego CA).

*Antibodies*

The following antibodies were used for IHC and ICC staining: SHH (1:250, Abcam Cat# ab53281, RRID:AB_882648), PTCH1 (1:250, Abcam Cat# ab53715, RRID:AB_882208), SMO (1:200, Abcam Cat# ab236465), Chromogranin A (1:100, Abcam Cat# ab254557, EPR22537-249), Synaptophysin (1:200, Cell Signaling Technology Cat# 9020, RRID:AB_2631095), GFAP (1:400, Agilent Cat# Z0334, RRID:AB_10013382), GFAP (1:1000, Abcam Cat# ab4674, RRID:AB_304558), Acetylated Tubulin (1:200, Sigma-Aldrich Cat# MABT868, RRID:AB_2819178), Ki-67 (1:200, Cell Signaling Technology Cat# 12202, RRID:AB_2620142), S100B (1:200, Abcam Cat# ab52642, RRID:AB_882426), $P75^{NTR}$ (1:1000, Abcam Cat# ab52987, RRID:AB_881682), and PLP1 (1:200, Abcam Cat# ab275751, RRID:AB_2915963). Secondary antibodies were used at 1:400 dilution: Donkey anti-Rabbit IgG AF488 (Thermo Fisher Scientific Cat# A-21206, RRID:AB_2535792), Donkey anti-Rabbit IgG AF594 (Thermo Fisher Scientific Cat# A-21207, RRID:AB_141637), Donkey anti-Rabbit IgG AF647 (Thermo Fisher Scientific Cat# A-31573, RRID:AB_2536183), Donkey anti-Mouse IgG AF488 (Thermo Fisher Scientific Cat# A-21202, RRID:AB_141607), Donkey anti-Mouse IgG AF594 (Thermo Fisher Scientific Cat# A-21203, RRID:AB_141633), Donkey anti-Mouse IgG AF647 (Thermo Fisher Scientific Cat# A-31571, RRID:AB_162542), Goat anti-Chicken IgY AF488 (Thermo Fisher Scientific Cat# A-11039, RRID:AB_2534096), Goat anti-Chicken IgY AF647 (Thermo Fisher Scientific Cat# A21449, RRID:AB_1500594), and Donkey anti-Goat IgG HRP (Abcam Cat# ab205723, RRID:AB_3065024).

**Cell Culture**

BON-1 and STC-1 cell lines were obtained from ATCC (Manassas, VA). BON-1 cells were grown in DMEM/Ham's F12 media supplemented with 10% FBS and penicillin-streptomycin (Corning Inc., Corning, NY). STC-1 cells were grown in DMEM supplemented with 10% FBS and penicillin-streptomycin (Corning Inc., Corning, NY). All cells were maintained at 37$^{o}$C and used at passage numbers 5-10. Cells were routinely tested every six months for mycoplasma contamination using the Universal Mycoplasma Testing kit (Cat #30-1012K, ATCC).

*Primary Enteric Glial Cell Isolation*

For each genotype condition and experiment, EGCs were isolated from 2 to 3 adult mice aged 6- to 12-months using a modified version of previously published protocols *(54,55)*. Following euthanasia of the mice, the proximal 4 cm of the duodenum including the antropyloric Brunner's glands, were removed and flushed with cold Dulbecco's PBS w/o Ca and Mg (DPBS). The intact intestine was gently pulled onto a wetted wooden Q-tip stick and the mesentery and blood vessels were carefully removed. The tissue was then cut into 2-3 mm segments and incubated in 25 mL of chelation buffer (2 mM EDTA, 10 mM HEPES, 2% FBS in DPBS) for 20 min at 37°C on a rotating shaker set to 200 rpm. The tissue solution was manually shaken 3 to 5 times before filtering through a 70-micron cell strainer. The unfiltered tissue was incubated a second time in fresh chelation buffer for 20 min at 37°C on a shaker. The solution was filtered again through a 70-micron strainer, and the unfiltered tissue was minced with scissors and incubated in up to 20 mL tissue digestion buffer (10 mM HEPS, 2% FBS, 1.5 mg/mL collagenase type 3, 40 µg/ml DNase I, prepared fresh in DPBS) on a shaker at 37°C for 15-20 min. The solution was filtered through a 40-micron strainer and centrifuged at 400 *x g* for 5 min at 4°C. The cells were resuspended in glial seeding media (DMEM/F12, 10 mM HEPES, 2 mM GlutaMAX, 10% FBS, 100 U penicillin-streptomycin, 1X amphotericin-gentamicin, G5 supplement, N-2, and Gem21 neural growth supplements) and plated into 6-wells coated with poly-L-lysine (Cat #3438-200-01, Cultrex, R&D Systems). After 16 to 24 h incubation at 37°C, the media was changed to glial growth media (same as seeding media except with 0.5% FBS instead of 10%) to inhibit the growth of fibroblasts and non-glial cells. For the generation of a pure $SOX10^{+}$ cell population, EGCs from *Sox10-CreER*$^{T2}$*-LSL-tdTomato* mice were treated with 2 µM 4-OHT for 48 h and then dissociated and sorted by FACS using the FACS Aria II. Cre negative cells were used to establish the gating parameters for sorting positive cells. Sorted cells were plated on poly-l-lysine coated plates and subcultured to generate an EGC line.

**Tumor Organoid Culture**

Human tumor organoid (tumoroid) lines were generated from surgically resected PNET tissues that were provided fresh or frozen in cryopreservation media containing 10% DMSO. Tumor tissues were diced into 2-3 mm segments and incubated in a series of EDTA incubations and digested with collagenase type 3 and DNase I, as described exactly in the previous methods section. The resulting tumor cell pellet was resuspended in ice-cold Matrigel without phenol red (Corning) and seeded into pre-warmed 24-well glass bottom or plastic tissue-culture plates. Tumoroids were subcultured in complete organoid media as published on previously *(6,34)*. Complete media consisted of Advanced DMEM/F12, B-27, N-2, GlutaMAX (2 mM), HEPES (10 mM), penicillin-streptomycin (100 U/mL), 1X amphotericin-gentamycin, N-acetylcysteine

(1 mM), mixed 1:1 v/v with filtered conditioned media (Advanced DMEM/F12 with 20% FBS) collected from LWRN cells (Wnt, R-spondin, Noggin expressing, ATCC). For PNET tumoroids, complete media was supplemented with ascorbic acid (284 μM), retinoic acid (100 nM), recombinant human FGF-2 (25 ng/mL), recombinant human EGF (50 ng/mL), recombinant human insulin (10 μg/mL), Y-27632 (10 μM), and A83-01 (500 nM). Media was supplemented with 10 μM SB202190, a selective p38 MAP kinase inhibitor and 10 nM nicotinamide (Tocris Bioscience, Bristol, UK) for the first day of passaging to prevent anoikis. Organoids were used for studies between 14 to 21 days post-seeding and between passage numbers 0 and 3.

Mouse tumoroid cultures were generated from fresh and cryopreserved tumor tissues using enzymatic digestion as described above. Due to the relatively small size of the tumors, the tissues were not subject to EDTA incubations and were directly minced into digestion buffer with collagenase and DNase I. The resulting PNET cells were resuspended as described above and grown in PNET organoid media. SI-NET tumoroids were grown in Complete Organoid Media supplemented with recombinant human EGF (50 ng/mL), Y-27632 (10 μM), and A83-01 (500 nM), CHIR 99021 (4 μM), and gastrin I (10 nM).

**In vitro Drug Studies**

The tumoroids and cells were treated as described in the subsequent methods sections. Drug compounds used in the studies include: human recombinant SHH N-terminal peptide (R&D Systems, Cat# 1845-GMP), SAG (Tocris, Cat# 4366), vismodegib (Tocris, Cat# 7710), sonidegib (Tocris, Cat# 7826), GANT61 (Tocris, Cat# 3191), and itraconazole (Tocris, Cat# 5981). *Men1* knockdown was accomplished using ON-TARGETplus Mouse *Men1* siRNA (Horizon Discovery, Cat# L-042675-01-0005, Layfayette, CO). A SMARTpool of four targeting and non-targeting small interfering RNAs were used. The cells were transfected with 25 nM siRNAs using Lipofectamine 3000 without the use of the P3000 reagent (Invitrogen) according to the manufacturer's instructions. Cells were analyzed after 72 h post *Men1* silencing or treatment with non-targeting siRNAs.

**EdU Fluorescence Assay**

The proliferation of tumoroid cells was determined used the Click-iT™ EdU Cell Proliferation Kit for Imaging and performed as described by the manufacturer (Thermo Fisher Scientific, Cat# C10337). Tumoroids were dissociated using a 25 G needle syringe and resuspended in Matrigel without phenol red. The cell suspension was seeded into 96-wells and incubated in growth media for 5 to 7 days prior to initiating treatment. Following treatment for 5 days, 1X EdU was added to the wells and allowed to incubate for 4 hours. The remaining steps were performed as described in the kit instructions to visualize EdU+ cells with an Alexa Fluor conjugated secondary antibody. The cells were counterstained with Hoescht dye prior to imaging on the ECHO Revolved micoscrope. The number of $EdU^{+}$ cells were quantified per image and normalized to the total number of cells as enumerated by Hoescht staining.

**Bromodeoxyuridine (BrdU) incorporation Assay**

BrdU incorporation was measured using a BrdU assay kit according to the manual provided (Cell Signaling Technologies, Cat# 6813). BON-1 and STC-1 cells were seeded at a density of 10,000 cells in 96-well plates. The cells were serum starved in 1% FBS containing media for 24 h prior to initiating treatment with the respective compounds. Treatments were initiated for tumoroids 2 to 3 days post seeding and in the absence of serum starvation. 1X BrdU was added to each well after 72 h and allow to incorporate overnight. Cells were fixed and denatured for 30 min, incubated with anti-BrdU antibody for 1 h at 24°C, washed, and incubated in HRP-conjugated secondary antibody for 30 min at 24°C. Samples were washed, incubated in TMB substrate, and the HRP reaction was stopped and visualized by measuring the optical density (OD) at 450 nm using the Gen5 Microplate Reader and Data Analysis Software (BioTec, Dorset, UK).

**TdTomato Fluorescence Growth Assay**

TdTomato$^+$ tumoroids were seeded at a 1:5 dilution into 24-well plates in Matrigel without phenol red. Two to three days after seeding, treatment was initiated by replacing the growth media with fresh media containing the respective drug compounds. Tumoroids were incubated for 72 h and then assessed for growth. Tumoroids were extracted from Matrigel by gently dissolving the mixture in cold PBS. Tumoroids were transferred to 96-well black plates in triplicate and centrifuged. TdTomato fluorescence was quantified using the CLARIOstar® Plus plate reader (BMG LabTech) by measuring fluorescence in the Texas Red channel and using a 5 mm by 5 mm scanning area for each well. Values were averaged across the triplicate wells and normalized to the DMSO vehicle control treatment group to account for seeding variations across independent experiments.

**Serum Hormone Enzyme-linked Immunosorbent Assays (ELISA)**

Serum hormone levels were evaluated exactly as reported previously *(20)*. Glucagon was measured with the Mouse Glucagon enzyme-linked immunosorbent assay (ELISA) Kit (Crystal Chem, #81518). Serum was diluted 1:10 (10 µL serum per well) and incubated with anti-mouse glucagon and HRP-secondary antibody overnight at 4 °C. Glucagon concentration was determined by HRP activity and measuring absorbance at 450 nm. Serum insulin levels were measured with the Ultra Sensitive Mouse Insulin ELISA Kit (Crystal Chem, #90080). Five µL serum (1:20) was incubated with anti-mouse insulin antibody overnight at 4°C. HRP-secondary activity was measured by absorbance. Serum GLP-1 levels were measured using the Mouse GLP-1 ELISA kit (Crystal Chem, #81508). Ten µL serum was incubated with anti-GLP-1 antibody overnight at 4°C and HRP-secondary activity was measured by absorbance. Serum gastrin was evaluated with the Gastrin Enzyme Immunoassay kit (Sigma, #RAB0200). Serum was diluted 1:4 (25 µL serum per well) and incubated with biotinylated gastrin peptide anti-gastrin-1 antibody overnight at 4 °C. Gastrin concentration was determined by HRP activity and absorbance at 450 nm. Prolactin levels were measuring using the Mouse Serum Prolactin ELISA Kit (Invitrogen, #EMPRL). Serum was diluted 1:20 (5 µL serum per well) and incubated with anti-mouse prolactin antibody overnight at 4 °C. HRP-secondary activity was evaluated by measuring absorbance at 450 nm. For all assays, each mouse serum sample was averaged across duplicate wells.

**Quantitative Polymerase Chain Reaction (qPCR)**

RNA was extracted from the tumoroids using the ReliaPrep miRNA Cell and Tissue Miniprep System following the manufacturer's instructions (Promega, Madison, Wisconsin). Between 0.1 to 1 µg of cDNA was synthesized using SuperScript VILO IV after treatment with ezDNAse to remove genomic DNA. Quantitative PCR was performed using PowerUp SYBR Green Master Mix (Invitrogen) on 5 to 10 ng of cDNA using the QuantStudio 3 Real-Time PCR System (Applied Biosystems, Waltham, MA) with the following cycling conditions: 2 minutes at

50 °C, 2 minutes at 95 °C, denaturing step for 1 second at 95 °C, extension and annealing for 1 minute at 60 °C, followed by a dissociation melt curve stage to confirm primer specificity. All forward and reverse primers were purchased as validated predesigned PrimeTime qPCR Assay primer sets used for SYBR Green dye (Integrated DNA Technologies, Coralville, IA). qPCR data was expressed as fold-change using the established $2^{-ddCt}$ method

**Western Blot Analysis**

Tissues were homogenized using a rotor-stator homogenizer in 250 to 500 µL of ice-cold RIPA buffer (Thermo Fisher Scientific) supplemented with 1X HALT protease and phosphatase inhibitor (Thermo Fisher Scientific). Tissues were homogenized on ice at 10,000 rpm over four 15 second intervals. For generating protein extracts from the tumoroids, tumoroids were scraped from wells and collected in ice-cold PBS. Tissue lysates were centrifuged for 15 min at 15,000 x g at 4°C and the supernatant was collected as the protein extract. Organoids were dissociated from the Matrigel (Corning, Corning, NY) by gently pipetting ten times and centrifuging for 5 min at 300 x g at 4°C. The organoid pellet was resuspended in 1 mL ice-cold PBS and then centrifuged for 5 min at 300 x g at 4°C. To generate whole cell extracts, the pellet was resuspended in 200 µL of ice-cold RIPA buffer (Thermo Fisher Scientific) supplemented with 1X HALT protease and phosphatase inhibitor (Thermo Fisher Scientific). Organoids were lysed by passing through a 20 G syringe ten times and vortexing. Following 40 min incubation on ice, organoid lysates were centrifuged for 15 min at 15,000 x g at 4°C and the supernatant was collected as the protein extract.

Protein extracts (10–15 µg) were prepared in reducing conditions with 1X SDS buffer with 5% β-mercaptoethanol and denatured by boiling at 95 °C for 5 minutes. Proteins were run on precast gradient gels (4-12% Bis-Tris, Invitrogen) in cold 1X MOPS Gel electrophorese buffer for 10 min at 80 V, then 90 min at 110 V. Proteins were transferred onto PVDF membranes using the iBlot 2 (Invitrogen). The membranes were blocked in 5% BSA in TBS with 0.05% Tween-20 (TBST) buffer for 1 h on a rocker at 24°C. Blots were incubated in primary antibodies that were diluted in 5% BSA-TBST at 4°C overnight on a rocker. Membranes were washed three times in TBST, then incubated in HRP-linked anti-mouse or anti-rabbit IgG antibody for 1 h at 24°C with gentle rocking (1:3000 dilution, Cell Signaling Technology). Membranes were washed in TBST and protein bands were visualized using the Pierce ECL detection system (Cat #32106, Thermo Fisher Scientific). For quantitation, films were scanned in gray scale and quantified using ImageJ (FIJI) software. Protein bands were identified by the expected molecular weights and quantified with the area selection tool using the inverted mean gray value (MGV) method. MGVs were subtracted from 255 (pixel density maximum) to obtain the inverted MGV, then background subtraction and normalization to respective loading controls was applied. Representative images of blots were enhanced using global brightness adjustments that were applied to the entire image.

*Antibodies*

Antibodies used for western blots are as follows: GLI1 (Cell Signaling Technology Cat# 3538, RRID:AB_1903989), GLI2 (Abcam, Cat #ab187386, OTI1F9), SHH (Abcam Cat# ab53281, RRID:AB_882648), PTCH1 (Abcam Cat# ab53715, RRID:AB_882208), SMO (Abcam Cat# ab236465), PAX6 (Cell Signaling Technology Cat# 60433, RRID:AB_2797599), Chromogranin A (Abcam Cat# ab254557, EPR22537-249), Synaptophysin (Cell Signaling Technology Cat# 9020, RRID:AB_2631095), GFAP (1:2000, Agilent Cat# Z0334,

RRID:AB_10013382), Pan-ERK1/2 (Cell Signaling Technology Cat# 9102, RRID:AB_330744), Phospho-ERK1/2 (T202/Y204) (Cell Signaling Technology Cat# 9101, RRID:AB_331646), Pan-AKT (Cell Signaling Technology Cat# 4685, RRID:AB_2225340), Phospho-AKT (S473) (Cell Signaling Technology Cat# 4060, RRID:AB_2315049), Menin (1:10000, Bethyl Laboratories Cat #A300-105A, RRID: AB_2143306), GAPDH (Cell Signaling Technology, Cat #5174, Clone D16H11, RRID: AB_10828810), and β-actin (Cell Signaling Technology Cat# 4970, RRID:AB_2223172).

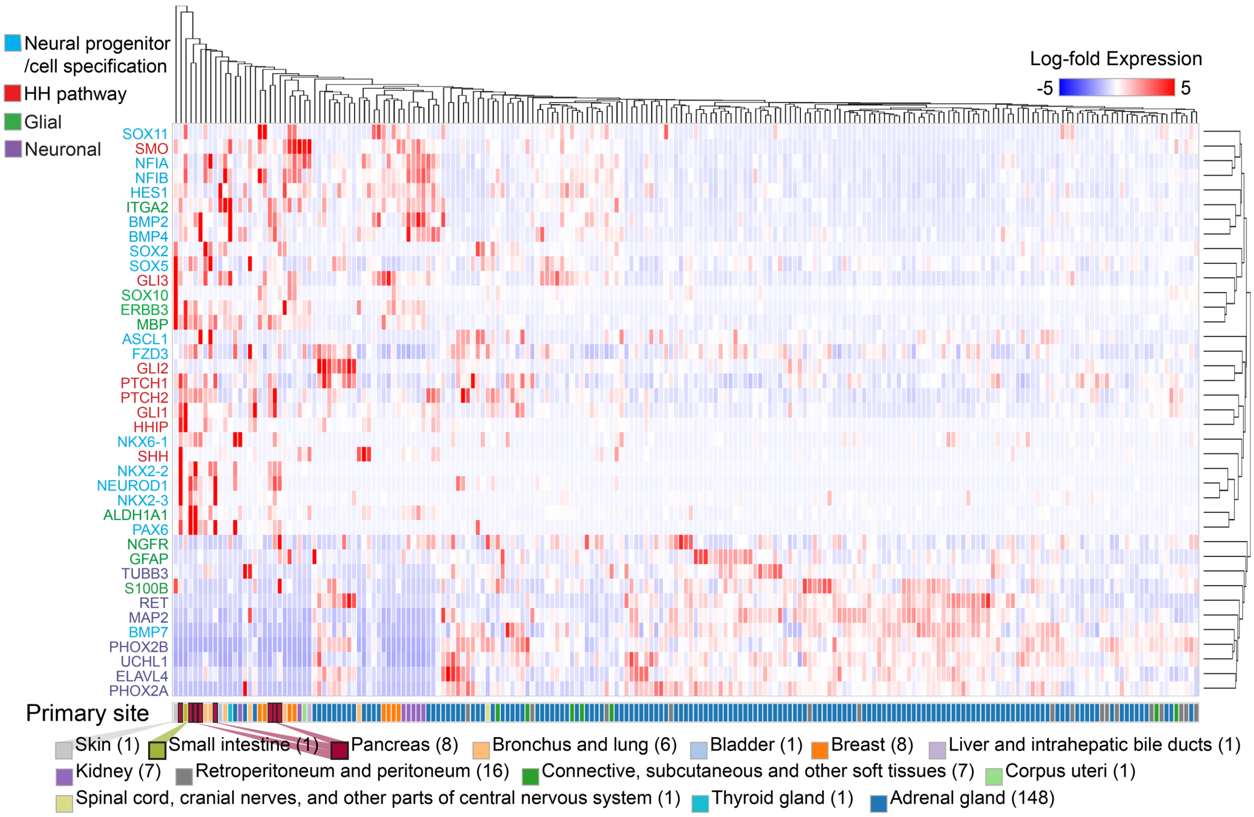


**Supplementary Figure 1.** Heatmap generated from whole transcriptome sequencing (WTS) data of TCGA-neuroendocrine cohort showing relative expression of transcripts mapped to neural progenitor cell specification, HH signaling pathway, and glial and neuronal cell types.

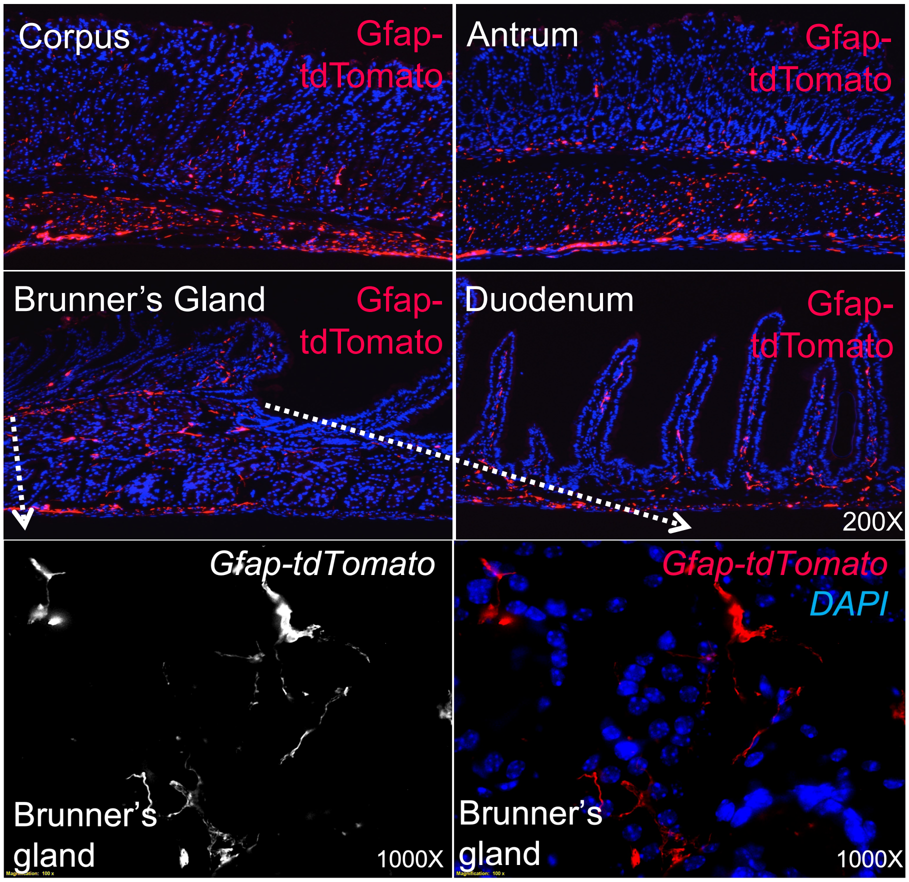


**Supplementary Figure 2.** TdTomato fluorescence of the gastric corpus, gastric antrum, Brunner's glands (BG), and duodenum of *GFAP-Cre; LSL-tdTomato* mice showing EGC distribution in the submucosa. Bottom panels show higher magnification images and highlight the typical glial morphology of $GFAP^{+}$ cells located in the Brunner's glands.

| Gene ID | Gene Symbol | Gene Name | Cytoband | Number of Cases Tested for Copy Number Alterations | Number of cases with CNV Gain | Percentage of cases with CNV Gain | Number of cases with CNV Loss | Percentage of cases with CNV Loss |
|---|---|---|---|---|---|---|---|---|
| ENSG00000128602 | SMO | smoothened, frizzled class receptor | 7q32.1 | 1407 | 774 | 55.01 | 19 | 1.35 |
| ENSG00000164690 | SHH | sonic hedgehog signaling molecule | 7q36.3 | 1407 | 758 | 53.87 | 31 | 2.2 |
| ENSG00000106571 | GLI3 | GLI family zinc finger 3 | 7p14.1 | 1407 | 708 | 50.32 | 11 | 0.78 |
| ENSG00000111087 | GLI1 | GLI family zinc finger 1 | 12q13.3 | 1407 | 120 | 8.53 | 85 | 6.04 |
| ENSG00000185920 | PTCH1 | patched 1 | 9q22.32 | 1407 | 80 | 5.69 | 108 | 7.68 |
| ENSG00000117425 | PTCH2 | patched 2 | 1p34.1 | 1407 | 72 | 5.12 | 275 | 19.55 |
| ENSG00000074047 | GLI2 | GLI family zinc finger 2 | 2q14.2 | 1407 | 33 | 2.35 | 44 | 3.13 |
| ENSG00000133895 | MEN1 | menin 1 | 11q13.1 | 1407 | 37 | 2.63 | 122 | 8.67 |

**Supplementary Table 1.** Prevalence of copy number variations (CNV) in hedgehog pathway genes for neuroendocrine cancers and cancers of glial cell origins in TCGA database.

| Gene ID | Gene Symbol | Gene Name | Cytoband | Number of Cases Tested for Copy Number Alterations | Number of cases with CNV Gain | Percentage of cases with CNV Gain | Number of cases with CNV Loss | Percentage of cases with CNV Loss |
|---|---|---|---|---|---|---|---|---|
| ENSG00000133895 | MEN1 | menin 1 | 11q13.1 | 165 | 37 | 22.42 | 122 | 73.94 |
| ENSG00000128602 | SMO | smoothened, frizzled class receptor | 7q32.1 | 165 | 79 | 47.88 | 3 | 1.82 |
| ENSG00000106571 | GLI3 | GLI family zinc finger 3 | 7p14.1 | 165 | 79 | 47.88 | 1 | 0.61 |
| ENSG00000117425 | PTCH2 | patched 2 | 1p34.1 | 165 | 17 | 10.3 | 32 | 19.39 |
| ENSG00000164690 | SHH | sonic hedgehog signaling molecule | 7q36.3 | 165 | 77 | 46.67 | 4 | 2.42 |
| ENSG00000111087 | GLI1 | GLI family zinc finger 1 | 12q13.3 | 165 | 22 | 13.33 | 23 | 13.94 |
| ENSG00000185920 | PTCH1 | patched 1 | 9q22.32 | 165 | 17 | 10.3 | 23 | 13.94 |
| ENSG00000074047 | GLI2 | GLI family zinc finger 2 | 2q14.2 | 165 | 7 | 4.24 | 16 | 9.7 |

**Supplementary Table 2.** Prevalence of copy number variations (CNV) in hedgehog pathway genes for *MEN1*-mutated cancers representing neuroendocrine or glial cell origins.
